# A multimodal arrayed Perturb-seq platform for dissecting microglial state dynamics and target identification

**DOI:** 10.64898/2026.11.16.767105

**Authors:** Linyang Ju, Matilda C. LaBranche, Stephanie Hromadka, Jingyao Li, Young Don Kwak, Xiaoping Chen, Jong Yu, Bindu Sukumaran, Kathleen A. Worringer, Christophe Freyre, Jack S. Hsiao

## Abstract

Microglia are highly dynamic immune cells of the central nervous system that can adopt diverse functional states in health and disease, making them both emerging therapeutic targets and challenging systems for target identification. Here, we developed TF-iMG, a human pluripotent stem cell-derived transcription factor-induced microglia model generated by inducible expression of SPI1 and CEBPA in Cas9-expressing stem cells. TF-iMG rapidly acquired microglia-like transcriptomic, morphological, inflammatory, and phagocytic features while remaining compatible with efficient lentiviral CRISPR knockout workflows that can be completed within one week.

We next established an arrayed multimodal Perturb-seq platform that integrates lentiviral CRISPR knockout, multiplexed single-cell RNA sequencing, cytokine profiling, and live-cell imaging. Single-cell analysis showed that TF-iMG recapitulates intrinsic microglial heterogeneity and captures homeostatic, activated, inflammatory, and proliferative signatures. Stimulation with IL1B and IL4 reproducibly induced distinct pro-inflammatory and anti-inflammatory states, respectively, providing a framework for perturbation-based interrogation of microglial state regulation. Targeting IL1B and IL4 pathway components validated the platform by attenuating expected stimulus-driven state shifts. Profiling of putative microglial regulators and Alzheimer’s disease-associated genes further revealed perturbation-specific transcriptional signatures, altered state occupancy, and multimodal phenotypes spanning cytokine release, morphology, and trajectory. Together, this platform enables scalable target identification in microglia.

## Introduction

Microglia are the resident immune cells of the central nervous system and are increasingly recognized as central regulators of brain homeostasis, injury responses, and neurodegenerative disease progression [1, 2]. In the healthy brain, microglia support synaptic remodeling, debris clearance, and tissue surveillance, whereas under pathological conditions they can adopt diverse reactive states that are either protective or detrimental depending on context [1, 3–6]. Increasing evidence has implicated these microglial state changes in the pathogenesis and progression of a wide range of neurological disorders, including Alzheimer’s disease (AD), amyotrophic lateral sclerosis (ALS), and other neuroinflammatory conditions [1, 2, 7–9].

CRISPR-based screening has emerged as a powerful framework for systematic gene-function studies, target discovery and validation [10–14]. In microglia, advances in human stem cell-derived models, high-throughput workflows, single-cell genomics, and CRISPR perturbation approaches have expanded the ability to study microglial state transitions in human *in vitro* systems with increasing resolution and scalability [15–25]. However, practical and biological challenges remain for microglia-focused CRISPR screening and assay development. For example, myeloid-lineage cells can be relatively refractory to efficient lentiviral transduction and genome editing, and conventional microglia differentiation workflows can require lengthy differentiation timelines or substantial optimization for scale, cryopreservation and CRISPR editing compatibility [26–28]. In addition, pooled CRISPR screens often rely on universal stress conditions and fitness-based endpoint readouts, which may not fully capture features that are important for microglia target prioritization, such as state heterogeneity, stimulus-dependent responses, or interactions with other neural cell types [6, 29]. For microglial target discovery, screening approaches that integrate multimodal transcriptomic, secreted, and phenotypic readouts and allow individualized experimental conditions in an arrayed format may therefore help distinguish whether a perturbation affects homeostasis, inflammatory activation, or protective responses, contextualized in disease-associated environments [29].

Perturb-seq, which combines CRISPR perturbation with single-cell RNA sequencing, provides an information-rich strategy for linking genetic perturbations to cell-state changes at single-cell resolution [10, 14, 30]. This approach has begun to be applied to human microglia models, as a downstream approach interrogating targets enriched from pooled CRISPR-screening [16, 23]. Yet despite its promise, comparatively few platforms have been tailored for microglia in an on-target arrayed screening format, combining single-cell transcriptomics and stimulus-defined microglial state mapping together with orthogonal functional readouts. For many target-discovery applications, a useful platform would support flexible experimental conditions, a robust and scalable microglia-like *in vitro* model, efficient genome editing, compatibility with disease-relevant stimuli, and integration of additional measurements such as cytokine release, morphology, and cell behavior [25, 29, 31].

Here, we developed such a platform by engineering a human pluripotent stem cell (hPSC)-derived transcription factor-induced microglia model, termed transcription factor-induced hPSC-derived microglia or TF-iMG, based on inducible overexpression of SPI1 and CEBPA in Cas9-expressing stem cells [15, 32]. This model rapidly generated microglia-like cells that exhibited canonical transcriptomic, morphological, inflammatory, and phagocytic features while also being compatible with efficient lentiviral CRISPR knockout workflows. Building on this system, we established an arrayed Perturb-seq platform that combined lentiviral CRISPR perturbation arrays, multiplexed single-cell RNA sequencing, cytokine profiling, and live-cell imaging in a unified experimental framework.

Using this platform, we first defined the intrinsic state landscape of TF-iMG and mapped stimulus-driven transitions across homeostatic, pro-inflammatory, and anti-inflammatory states. We then applied arrayed Perturb-seq to validate pathway regulators downstream of IL1B and IL4 and to profile putative microglial state regulators and disease-associated genes, revealing perturbation-specific states and disease-associated signatures. Together, this study established a rapid, scalable, and multimodal screening framework for microglia target identification and provides a practical approach for dissecting gene-dependent state dynamics in human microglia-like cells.

## Results

### hPSC-derived engineered TF-iMG exhibited microglia-like characteristics and is compatible with rapid CRISPR screening

To enable scalable target identification in human microglia, we sought to establish a stem cell-derived microglia model that combines rapid differentiation, robust microglial phenotypes, and compatibility with CRISPR-based screening. Inspired by previous transcription factor-driven microglia differentiation strategies [15], we engineered hPSCs to inducibly express the microglia-essential transcription factors SPI1 and CEBPA (Fig. 1A). In contrast to earlier implementations, we used a PiggyBac transposon strategy to stably integrate the inducible system in a constitutive Cas9 expressing H1-hPSC line [15, 32] (Fig. S1A). We refer to this model as TF-iMG, designed specifically for high-throughput CRISPR and multimodal screening applications.

**Fig. 1.**
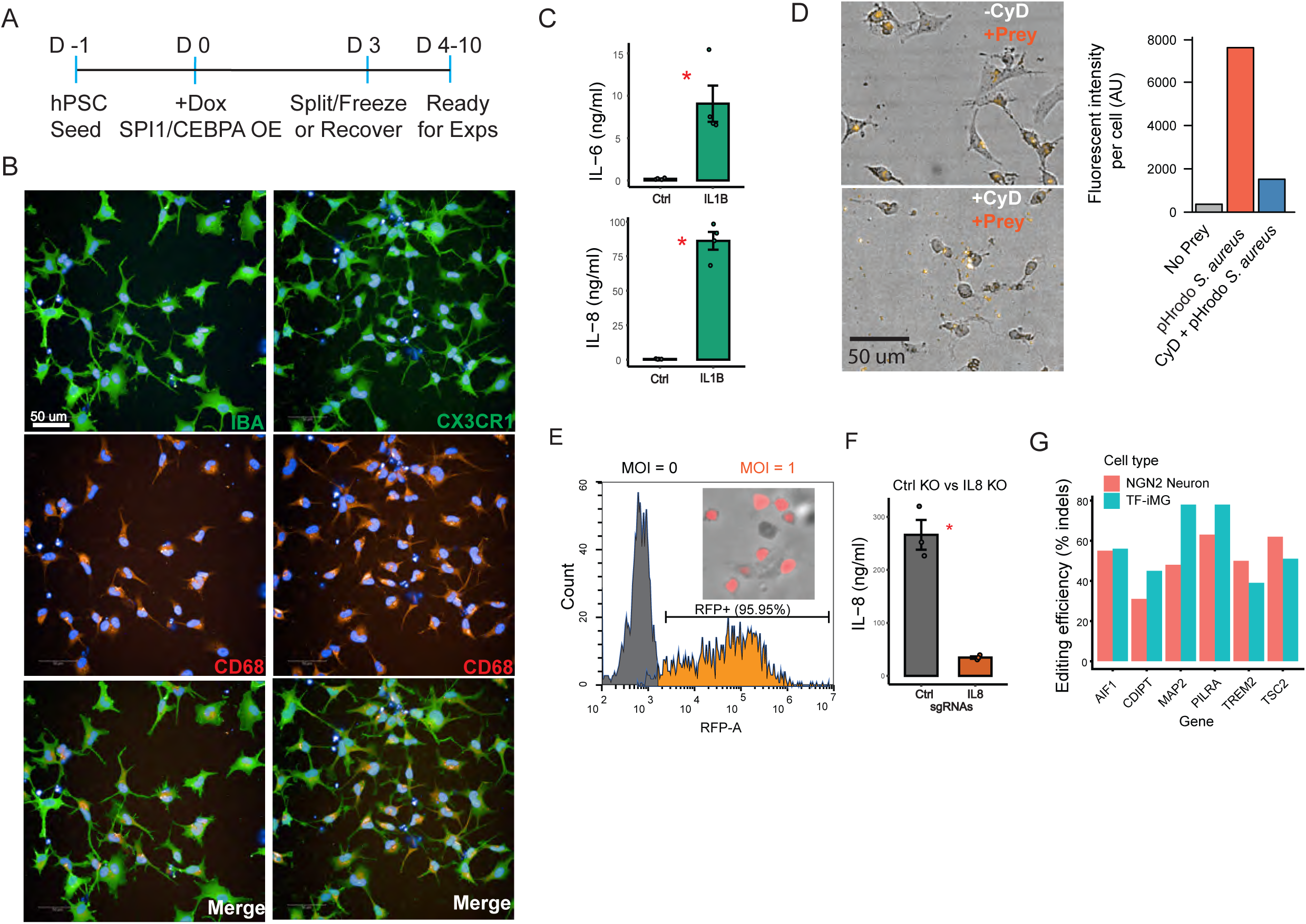
Generation and phenotypic characterization of the TF-iMG microglia model for CRISPR and multimodal screening. **Fig. 1A**. 4-day workflow of doxycycline (DOX)-induced TF-iMG differentiation from hPSCs. **Fig. 1B**. Representative immunofluorescence images of TF-iMG co-stained using IBA (left) and CX3CR1 (right) in green, CD68 in red, and DAPI in blue. **Fig. 1C**. Bar plots showing IL6 (top) and IL8 (bottom) release from TF-iMG following stimulation with IL1B, compared with BSA-treated controls. N = 4, *p < 0.1, Mann-Whitney U test. **Fig. 1D**. Phagocytic activity of TF-iMG assessed using pHrodo-conjugated *S. aureus* bioparticles. Left, representative images of TF-iMG at 18 h after bioparticle addition (top); the TF-iMG showed reduced uptake following cytochalasin D treatment (bottom). Right, bar plot of pHrodo fluorescence intensity per cell. **Fig. 1E**. Flow cytometry histograms of TF-iMG 3 days after lentiviral treatment at MOI = 0 and MOI = 1, with a representative image overlaying brightfield and RFP signals. **Fig. 1F**. Bar plots showing IL8 protein levels post 24h IL1B stimulation after 5-day CRISPR knockout of IL8 or non-targeting control knockout in TF-iMG. N = 3, *p < 0.1, Mann-Whitney U test. Fig. 1G. Bar plots showing CRISPR editing efficiency (indel percentage) after 6-day low-dose lentiviral treatment.

Consistent with the Chen et al. 2021 study, TF-iMG rapidly acquired microglia-like characteristics as early as 3 days after DOX induction, and by day 4 they showed pronounced ramified morphologies together with high expression of canonical microglial markers, including IBA1, CX3CR1, CD68, P2RY12, and CD11B validated by immunofluorescence staining or flow cytometry[15, 33–36] (Fig. 1B, Fig. S1B). To further characterize TF-iMG, we generated bulk RNA-seq and compared with multiple previously reported transcriptomic datasets, including *in vivo* postnatal and adult microglia, and conventional hPSC-derived microglia-like cells (iMG) and their integration into cerebral organoids (oMG) [37] (Fig. S1C). At 5 days after DOX induction, TF-iMG exhibited transcriptomic similarity to these reference datasets (Spearman coef = 0.607 – 0.747; Fig. S1C), further confirming their microglial identity.

We next assessed whether TF-iMG recapitulated key microglial functions relevant to disease modeling. Given that inflammatory responsiveness is a defining feature of microglial biology, we measured cytokine release following stimulation with IL1B [33] (Fig. 1C). TF-iMG responded to pro-inflammatory IL1B, with significant induction of pro-inflammatory cytokines such as IL6 and IL8 [38–41] (Fig. 1C). Additionally, TF-iMG efficiently internalized pHrodo-labeled *S. aureus* bioparticles; internalization was markedly suppressed by the addition of cytochalasin D (CyD) [15, 33] (Fig. 1D).

A central motivation for developing TF-iMG was to generate a system suitable for efficient CRISPR perturbation, addressing the well-recognized difficulty of transducing and editing mature myeloid-lineage cells [42]. To address this, we implemented lentiviral sgRNA delivery in combination with Vpx virus-like particles and achieved high lentiviral integration rates within 3 days [43] (>95% at target MOI = 1; Fig. 1E, FigS1A). Consistent with effective functional perturbation, IL8 sgRNAs (MOI =1) substantially reduced cytokine protein levels within 4 days, with IL8 levels decreased by 91%, compared with non-targeting sgRNA controls (Fig. 1F). To further assess editing efficiency in TF-iMG, we performed a parallel comparison with NGN2 neurons at day 6 post-transduction. Across the tested genes, editing efficiencies ranged from 39% to 78% and were overall comparable between TF-iMG and NGN2 neurons (Fig. 1G). Taken together, these results established TF-iMG as a rapid, scalable, and screening-compatible human microglia model that recapitulated key molecular and functional hallmarks of microglia.

### Single-cell transcriptomics revealed intrinsic TF-iMG state dynamics and stimulus-driven polarization into disease-relevant inflammatory states

Given that microglia occupy and can dynamically transition to a diverse landscape of transcriptomic states, we next asked whether TF-iMG could capture and recapitulate this complexity at single-cell resolution [16, 43–47]. We performed a pilot scRNA-seq experiment on untreated TF-iMG using a split-pool combinatorial barcoding workflow (Parse Biosciences) compatible with downstream high-throughput screening [14, 48, 49]. Pan-microglia genes, including canonical CSF1R and over-expressed SPI1, were broadly expressed across the population, confirming the overall microglial identity of the model [15, 16, 34, 43, 46] (Fig. 2A, 2B). Unsupervised clustering revealed five transcriptionally distinct subpopulations characterized by key markers corresponding to homeostatic (CX3CR1), activated (LRRK2), inflammatory (LYZ, SERPINB2), and proliferative (TOP2A) transcriptional states [16, 43–47] (Fig. 2A, 2B; Tables S1, S2). These findings indicated that TF-iMG were not locked into a single endpoint state but instead recapitulated intrinsic microglial heterogeneity similar to previous reports [16, 43–47].

**Fig. 2.**
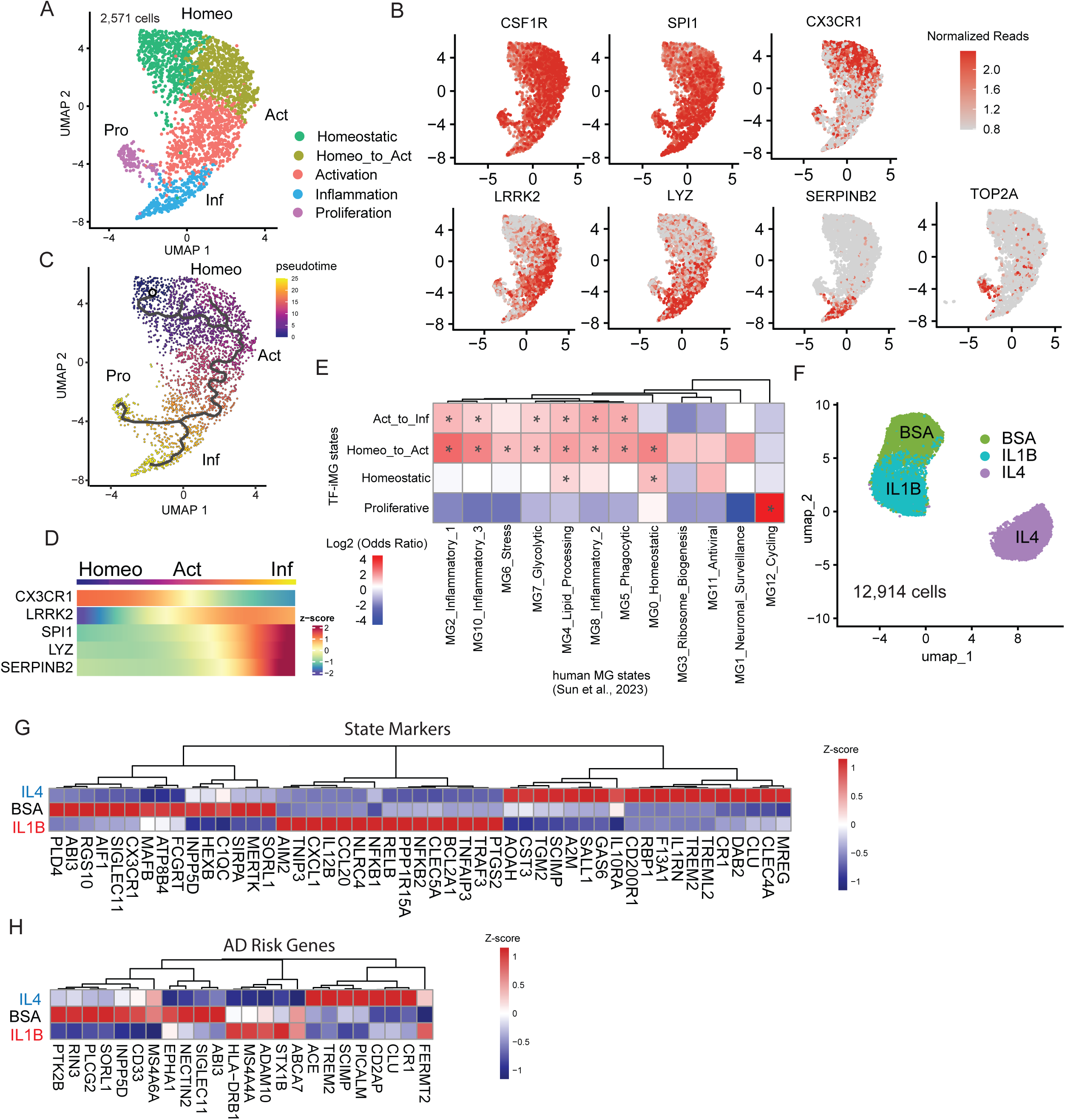
Single-cell transcriptomics reveals TF-iMG dynamics, human microglia signatures, and disease-relevant pro- and anti-inflammatory state shifts induced by IL1B and IL4. **Fig. 2A**. UMAP of day-5 TF-iMG scRNA-seq data. **Fig. 2B**. Feature plots showing expression of pan-microglia and state-associated marker genes in TF-iMG. Pan-microglia markers: CSF1R, SPI1; homeostatic: CX3CR1; activated: LRRK2; inflammatory: LYZ, SERPINB2; proliferative: TOP2A. **Fig. 2C**. UMAP colored by pseudotime trajectory illustrating TF-iMG state dynamics along a homeostatic-to-inflammatory axis. **Fig. 2D**. Heatmap of microglial state marker expression across the pseudotime trajectory in TF-iMG (z-scores). **Fig. 2E**. Enrichment of human microglia state signatures (identified in Sun et al. 2023) in corresponding TF-iMG states, shown as log2-transformed odds ratios. *p < 0.1E-04, Fisher’s exact test adjusted using Benjamini-Hochberg procedure. **Fig. 2F**. UMAP of day-5 TF-iMG profiled 24 h after treatment with BSA, IL1B, or IL4. **Fig. 2G**. Heatmap showing upregulated homeostatic, pro-inflammatory, and anti-inflammatory microglia markers following BSA, IL1B, and IL4 treatment in TF-iMG (z-scores). **Fig. 2H**. Heatmap of AD risk genes showing dynamic expression patterns associated with BSA, IL1B, or IL4 treatment in TF-iMG (z-scores).

To further define these state relationships, we performed pseudo-time trajectory analysis [50, 51]. This revealed a continuous trajectory spanning homeostatic, activated, and inflammatory states, here summarized as a “Homeo-Act-Inf” axis (Fig. 2C). Expression of key state markers changed progressively along this trajectory, supporting the interpretation that TF-iMG captures ongoing and ordered state transitions rather than disconnected clusters (Fig. 2D, Fig. S2A). To assess the biological relevance of these states, we compared TF-iMG cluster markers with the human microglia atlas reported by Sun et al. 2023 [46] (Fig. 2E; Table S2). Reciprocal enrichment analysis showed that TF-iMG states significantly overlapped with human *in vivo* microglia signatures, including homeostatic (MG0 Homeostatic), proliferative (MG12 Cycling), and multiple activated or inflammatory states such as MG2, 8, and 10 inflammatory, MG4 Lipid Processing, and MG5 Phagocytic (Fig. 2E). Thus, TF-iMG reproduces not only generic microglia marker expression, but also a spectrum of transcriptional states observed in human microglia.

We next asked whether TF-iMG could transition into defined inflammatory states relevant to disease modeling. We selected IL1B and IL4 as prototypical stimuli representing pro-inflammatory and anti-inflammatory polarization, respectively, and profiled TF-iMG 24 hours after treatment alongside BSA-treated controls [40, 46, 52–56]. Integrated single-cell analysis showed that both IL1B and IL4 induced distinct and separable transcriptional states relative to control cells (Fig. 2F). Differential expression analysis highlighted markers generally defining or associated with homeostatic, pro-inflammatory, and anti-inflammatory states, such as CX3CR1 and INPP5D in homeostatic cells, IL12B and NFKB1 in IL1B-treated cells, CD200R1 and TREM2 in IL4-treated cells [1, 43, 46, 57–64] (Fig.2G; Table S2). These data defined a tractable *in vitro* state framework spanning homeostatic, pro-inflammatory, and anti-inflammatory microglial states essential for downstream perturbation studies.

Notably, these stimulus-defined states also displayed distinct expression patterns of genes linked to Alzheimer’s disease risk (Fig. 2H). For example, ABI3 was elevated in the homeostatic state, MS4A4A in the pro-inflammatory state, and CLU in the anti-inflammatory [65–67] (Fig. 2H). This supported the disease relevance of the TF-iMG state space and suggested that defined microglial activation states may differentially engage neurodegeneration-associated pathways.

To extend the platform beyond canonical inflammatory cues, we also profiled TF-iMG treated with astrocyte-conditioned medium (ACM) [68]. ACM induced an additional transcriptional state distinct from both IL1B- and IL4-polarized cells (Fig. S2B). This state was marked by upregulation of genes associated with microglial activation and metabolic adaptation, including IL1R2, IL18R1, CD38, CD163, TLR2, and TLR8 [69–72] (Fig. S2C; Table S2). Gene ontology analysis further highlighted terms related to chronic inflammatory responses, acute-phase response, interleukin-1 production, and lipid-associated metabolic signatures (Fig. S2D; Table S3). Together, these results showed that TF-iMG supported both intrinsic and stimulus-induced microglial state transitions and provided a biologically meaningful reference map for downstream perturbation studies.

### Arrayed Perturb-seq with multimodal readouts validated pathway-level regulators of IL1B- and IL4-driven microglial state transitions

Given the compatibility of TF-iMG with rapid CRISPR editing and its reproducible stimulus-driven state transitions, we next developed an arrayed Perturb-seq workflow to identify perturbations that would drive shifts or changes in microglial transcription states. To enable multi-dimensional assessment of perturbation effect, we combined CRISPR perturbation with multiplexed scRNA-seq, cytokine profiling and live-cell imaging as parallel orthogonal readouts (Fig. 3A). As a first proof-of-concept application, we focused on genes within the IL1B and IL4 signaling pathways, reasoning that perturbation of these pathway components should attenuate the corresponding pro- or anti-inflammatory state transitions. The target panel included IL1R1, IL1R2, CXCL8, NFKB1, and TRAF6 for the IL1B pathway; IL4R, JAK1, IL2RG, and STAT6 for the IL4 pathway; a non-targeting control; and TGFBR2, which has been previously reported to disrupt microglial homeostasis, as a positive control [16, 34, 73, 74] (Fig. 3B).

**Fig. 3.**
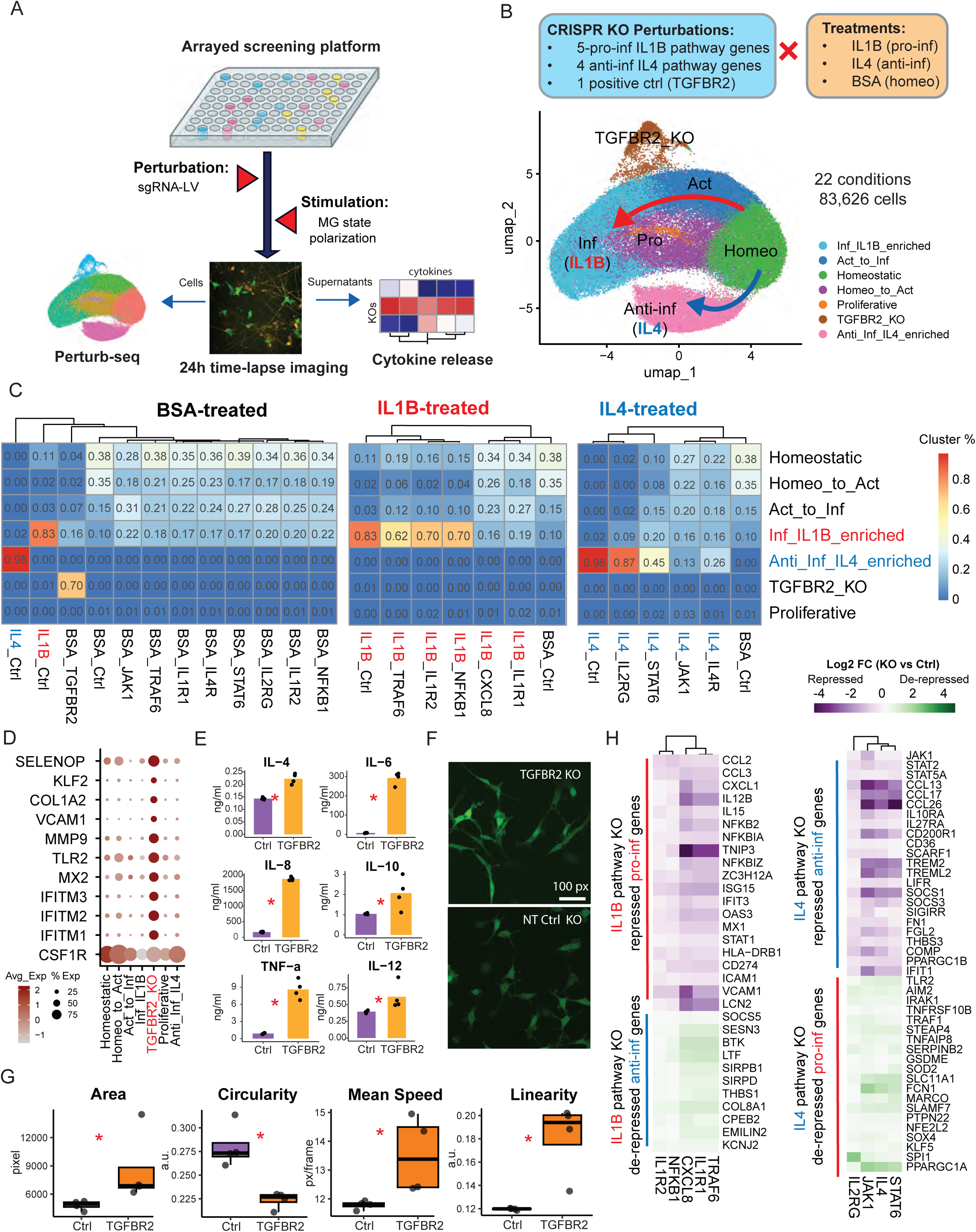
Arrayed Perturb-seq in TF-iMG targeting regulators of IL1B- and IL4-driven state transitions. **Fig. 3A**. Schematic overview of the arrayed Perturb-seq workflow in TF-iMG with integrated time-lapse imaging and cytokine profiling. **Fig. 3B**. Experimental design (top), showing 22 treatment-by-perturbation conditions, and integrated UMAP (bottom) of TF-iMG arrayed Perturb-seq targeting IL1B and IL4 pathway regulators together with control perturbations. **Fig. 3C**. Heatmap showing the proportion of cells assigned to each microglia state across perturbation conditions, grouped by BSA (left), IL1B (middle), and IL4 (right) treatment. **Fig. 3D**. Dot plot of top marker genes selectively enriched in the TGFBR2 knockout cluster in TF-iMG. **Fig. 3E**. Bar plots showing significant upregulation of pro-inflammatory cytokines (IL6, IL8, TNF-alpha, and IL12) as well as anti-inflammatory cytokines (IL4 and IL10) following TGFBR2 knockout compared with non-targeting control knockout. N = 4, *p < 0.1, Mann-Whitney U test. **Fig. 3F**. Representative images of TGFBR2 knockout TF-iMG (top) and non-targeting control knockout TF-iMG (bottom) at day 9, corresponding to 5 days after lentiviral transduction. **Fig. 3G**. Box plots showing significant changes in cell morphology parameters (area and circularity) and trajectory parameters (mean speed and linearity) quantified from 24-h time-lapse imaging of TGFBR2 knockout versus non-targeting control TF-iMG. N = 4, *p < 0.1, Mann-Whitney U test. **Fig. 3H**. Heatmap of top IL1B-responsive (left) and IL4-responsive (right) genes that are systematically repressed or de-repressed following perturbation of IL1B or IL4 pathway components in TF-iMG (z-scores).

In the arrayed workflow, day 4 TF-iMG cells expressing constitutive GFP were transduced with lentiviral constructs carrying three sgRNAs per target at MOI = 1 for 5 days, followed by stimulation with IL1B, IL4, or BSA control on day 9. For selected conditions, 24-hour time-lapse imaging and cytokine profiling were performed in parallel (Fig. 3A). Across 22 conditions representing 11 targets under different stimulation contexts, we recovered 83,626 cells for scRNA-seq analysis (Fig. 3B). Integrated analysis confirmed that the key transcriptional features established in earlier experiments were preserved in this perturbation setting: TF-iMG retained intrinsic homeostatic-to-inflammatory heterogeneity, and IL1B and IL4 continued to induce robust pro- and anti-inflammatory state shifts (Fig. 3B, Fig. S3A; Tables S1, S2).

We first examined the positive control perturbation TGFBR2. As expected from previous reports, TGFBR2 knockout produced a distinct transcriptional cluster not observed in other conditions [16] (Fig. 3B, 3C; Table S1). Although pan-microglia markers remained expressed, TGFBR2-deficient cells upregulated a set of genes including IFITM1, IFITM2, IFITM3, MX2, and MMP9, consistent with a dysregulated and activated microglial state [75–77] (Fig. 3D; Table S2). Pathway enrichment analysis identified TGF-beta signaling; in addition, ECM-receptor interaction and complement and coagulation cascades were major signatures in this cluster, which were also closely associated with TGF-beta signaling [34, 56, 78] (Fig. S3B; Table S3). Importantly, this transcriptomic dysregulation was mirrored in orthogonal assays: TGFBR2 knockout strongly increased release of multiple cytokines, including IL4, IL6, IL8, IL10, IL12, and TNF-alpha (Fig. 3E), and live imaging revealed altered morphology and motility, with cells becoming larger, less circular, and more motile and directed than control cells (Fig. 3F, 3G). These results confirmed that the platform could detect strong perturbation-induced deviations in microglial state across transcriptomic, secreted, and phenotypic dimensions.

We next evaluated perturbations of the IL1B and IL4 signaling pathways. Across these conditions, we observed that most perturbations shifted cells in the expected direction, reducing the extent of the induced inflammatory transition even when the effect was partial rather than absolute (Fig. 3C). In particular, knockout of IL1R1 and CXCL8 attenuated the IL1B-driven pro-inflammatory transition. The IL1B_enriched population dropped from 83% to 19% and 16% respectively. Knockout of JAK1, IL4R, and STAT6 blocked the IL4-driven anti-inflammatory shift. The IL4_enriched population dropped from 98% to 13%, 26%, and 45% respectively (Fig. 3C, Fig. S3C; Table S1). Although not every perturbation produced a complete redistribution of cell states, pathway-level effects were consistently detectable at the gene expression level, where we observed that robust IL1B/IL4-responding genes were consistently repressed or de-repressed after perturbation across all targets (Fig. 3H; Table S4). For example, blockades of IL-1β signaling suppressed IL-1β-responsive genes such as CCL2 and de-repressed anti-inflammatory modulators such as LTF. In contrast, perturbation of IL-4 signaling reduced IL-4-responsive genes such as CCL17 and up-regulated pro-inflammatory pattern-recognition receptors such as TLR2 [79–84] (Fig. 3H; Table S4). Cytokine measurements provided further orthogonal support: IL1R1 and CXCL8 knockout reduced release of pro-inflammatory cytokines including IFN-gamma, IL6, and IL8 under IL1B stimulation (Fig. S3D). Together, these findings validated the arrayed Perturb-seq workflow as a robust platform for linking defined gene perturbations to stimulus-specific microglial state changes.

### Multimodal arrayed Perturb-seq identified perturbation-specific and disease-relevant signatures of microglial state regulators

To further test the discovery potential of the platform, we performed a second pilot arrayed Perturb-seq experiment focused on microglial state regulators and disease-associated genes. We selected IRF8 and HIF1A as putative TF regulators of microglial activation state, and TREM2 and ABI3 as disease-relevant microglia-enriched genes, and profiled these perturbations alongside a non-targeting control under BSA, IL1B, and IL4 treatment [46, 65, 85–87] (Fig. 4A). In this experiment, we integrated a full multimodal readout strategy comprising scRNA-seq, 11 cytokine measurements (Table S7), and 18 quantitative imaging-derived phenotypic features (Table S8).

**Fig. 4.**
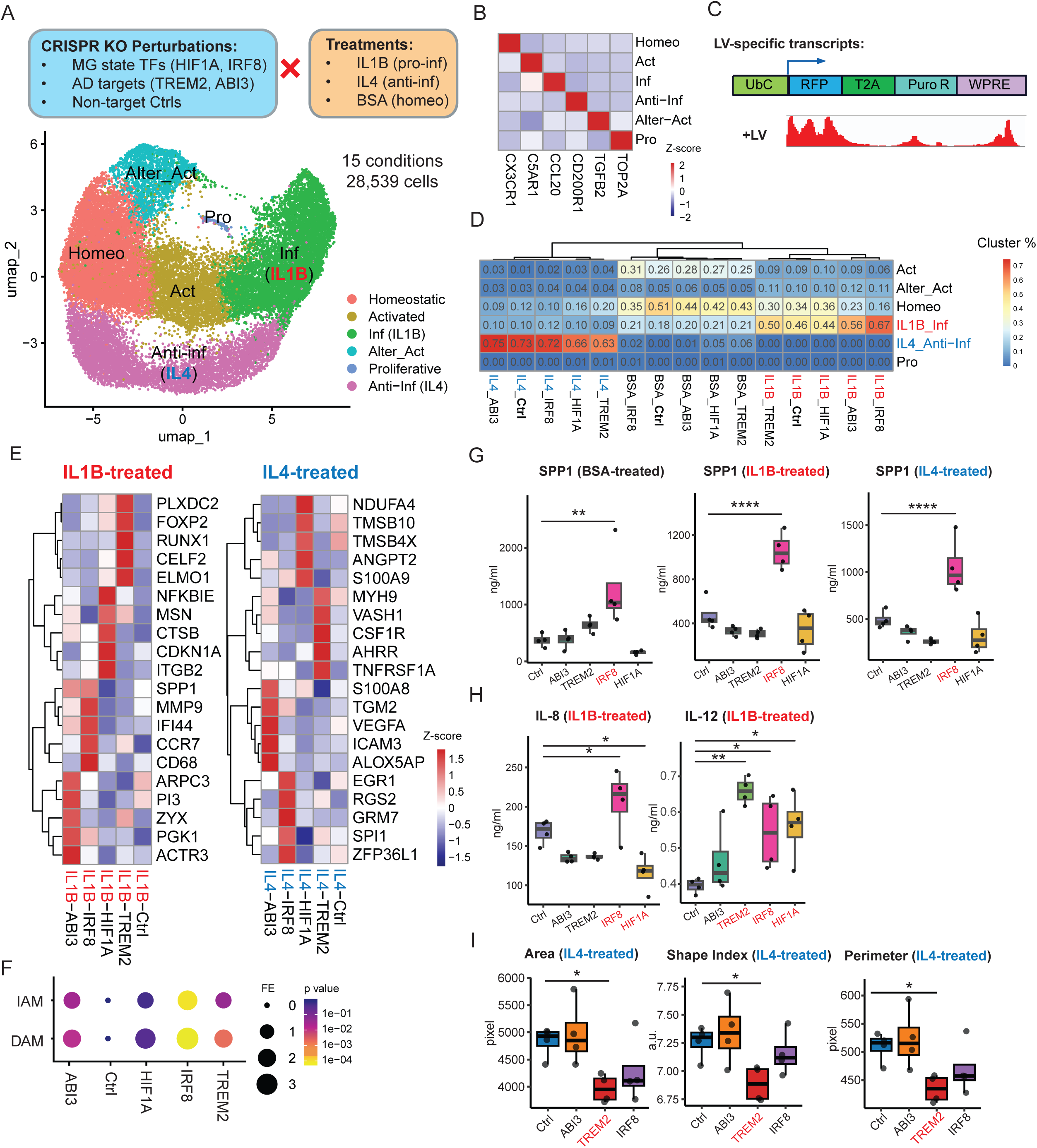
Multimodal arrayed Perturb-seq identifies disease-relevant signatures of diverse microglial state regulators. **Fig. 4A**. Experimental design (top), showing 15 treatment-by-perturbation conditions, and integrated UMAP (bottom) of TF-iMG arrayed Perturb-seq targeting transcription factors and disease-relevant genes. **Fig. 4B**. Heatmap showing consistent microglia state marker expression across the dataset (z-scores). **Fig. 4C**. Genome browser tracks showing lentiviral marker transcripts aligned to the lentiviral reference genome in lentivirus-treated cells (top) and no-lentivirus controls (bottom). **Fig. 4D**. Heatmap showing the proportion of cells assigned to each microglia state across perturbation conditions, grouped by IL4 (left), BSA (middle), and IL1B (right) treatment. **Fig. 4E**. Heatmap of curated knockout-exclusive upregulated genes following IL1B (left) or IL4 (right) polarization (z-scores). **Fig. 4F**. Dot plot showing fold enrichment (FE) of DAM and IAM signatures among knockout-specific upregulated genes under IL1B stimulation, assessed by Fisher’s exact test. **Fig. 4G**. Box plots showing SPP1 release across all five perturbations under BSA (left), IL1B (middle), and IL4 (right) treatment conditions. N =4. Statistical significance for Fig. 4G-I and Fig. S4B-E was determined by ANOVA followed by Dunn’s test comparing each perturbation with the non-targeting control. *p < 0.1, **p < 0.01, ***p < 0.001, ****p<0.0001. **Fig. 4H**. Box plots showing significantly altered cytokine release across all five perturbations under IL1B polarization. N = 4. **Fig. 4I**. Box plots showing significantly altered cell morphology parameters under IL4 polarization quantified from 24-h time-lapse imaging across all perturbations except HIF1A. N = 4.

Across 15 conditions, we recovered 28,539 cells and again observed a consistent microglial state landscape extending across homeostatic, activated, pro-inflammatory, anti-inflammatory, and proliferative compartments, marked by genes such as CX3CR1, C5AR1, CCL20, CD200R1, and TOP2A (Fig. 4A, 4B; Tables S1 and S2). To directly assess perturbation capture, we built a custom reference genome that included lentiviral transcript sequences. Using this reference, we detected lentiviral marker transcripts in 75% of cells overall, with detection rates across conditions ranging from 64.7% to 87.1% (Fig. 4C, Fig. S4A). These data supported the technical robustness of the perturbation workflow.

Unlike pathway-level perturbations in the IL1B/IL4 validation experiment, knockout of IRF8, TREM2, ABI3, or HIF1A did not result in a strong binary modulation of inflammatory responses. Instead, these perturbations more subtly reshaped the distribution of cells across stimulus-defined states (Fig. 4D; Table S1). This was particularly evident under IL1B stimulation, where IRF8 knockout increased the proportion of cells occupying the inflammatory state relative to the respective non-targeting control gRNA-treated condition (67% vs 46%), representing the strongest pro-inflammatory bias among all tested perturbations [87]. In contrast, under IL4 treatment, TREM2 knockout produced a slight shift toward the anti-inflammatory cluster (63% vs 73%), consistent with a role for TREM2 in supporting protective microglial responses [58, 88] (Fig. 4D).

To identify perturbation-specific transcriptional consequences, we curated genes exclusively upregulated in each knockout condition under IL1B or IL4 polarization. Each perturbation displayed a distinct transcriptional signature containing genes linked to the perturbed pathway or disease context (Fig. 4E; Table S4). For example, SPP1, a disease-associated microglia (DAM) marker, was induced by IRF8 perturbation; PLXDC2 was elevated in TREM2-deficient cells; and PI3 and CDKN1A responded to ABI3 and HIF1A perturbation, respectively [65, 89–91] (Fig. 4E; Table S4). To further place these transcriptional effects in disease context, we evaluated enrichment of DAM and inflammation-associated microglia (IAM) signatures in KO-specific upregulated genes under IL1B stimulation [86, 92, 93] (Fig. 4F; Table S5). All targeted perturbations showed enrichment for DAM and IAM signatures relative to control-biased genes, with IRF8 knockout showing the strongest enrichment, in agreement with its pronounced inflammatory state shift (Fig. 4F).

The orthogonal readouts reinforced these transcriptomic findings. Because SPP1 emerged as a key marker in the scRNA-seq data and is a known IRF8-regulated factor, we incorporated SPP1 into the cytokine panel. Consistent with the transcriptional analysis, IRF8 knockout significantly increased SPP1 secretion across all treatment conditions (Fig. 4G) and also elevated pro-inflammatory cytokines such as IL8 and IL12 under IL1B stimulation (Fig. 4H). TREM2 knockout also produced a dysregulated cytokine profile, including reduced IL6 and IL10 under IL4 treatment (Fig. S4B) and altered balances of IL12, and IL13 under IL1B treatment (Fig. 4H, Fig. S4B), consistent with disruption of protective microglial signaling [94]. Additional perturbations including HIF1A and ABI3 likewise yielded selective cytokine abnormalities, underscoring the utility of parallel secretome profiling for interpreting transcriptomic hits (Fig. 4H, Fig. S4B, S4C).

Live-cell imaging provided a further layer of phenotypic resolution. Under IL4 polarization, TREM2 knockout selectively altered morphology-related features including cell area, shape index, and perimeter, producing a smaller and more rounded phenotype suggestive of impaired acquisition of the IL4-associated protective state [58] (Fig. 4I). By contrast, IRF8 and ABI3 knockout more strongly affected trajectory-related parameters such as linearity and confinement ratio (Fig. S4D). IRF8 knockout caused cells to wander more off course only under IL1B stimulation, whereas ABI3 knockout impaired their ability to travel in a direct path under both IL1B and IL4 conditions (Fig. S4D, S4E). Although the mechanistic basis of these migration-associated phenotypes will require further validation, these results showed that multimodal analysis can reveal functional dimensions of microglial dysregulation that are were not fully captured by transcriptomics alone.

Together, these findings demonstrated that the TF-iMG arrayed Perturb-seq platform can be leveraged not only for pathway validation but also to uncover disease-relevant regulators that modulate microglial state in complex ways. By integrating transcriptomic, cytokine, and time-lapse imaging readouts, the platform resolved graded state shifts, perturbation-specific downstream gene signatures, and convergent phenotypic consequences across molecular, secreted, and imaging-based modalities.

## Discussion

### A rapid and screening-compatible human microglia model

Microglia are increasingly viewed as central regulators of neurological disease, but their highly dynamic and heterogeneous biology makes them challenging when trying to identify or validate a potential target and establish therapeutic strategies to modulate that target [95]. In this study, we developed a rapid, scalable, and multimodal screening platform designed specifically to address that challenge. By combining an engineered human stem cell-derived microglia model with arrayed CRISPR perturbation, single-cell transcriptomics, cytokine profiling, and live imaging, we established an experimentally flexible framework for dissecting how genetic perturbations reshape microglial states.

We have developed a TF-iMG model equipped with Cas9-enabled editing. Compared with more conventional stem cell-derived microglia differentiation approaches [33], TF-iMG offered several practical advantages that are particularly important for screening applications, including rapid differentiation, compatibility with cryopreservation, stable Cas9 expression, and efficient lentiviral CRISPR editing. However, TF-iMG did not respond as expected to some perturbations, such as LPS, commonly used to induce a response from microglia differentiated using conventional methods; furthermore, maintaining their long-term survival under current experimental conditions proved challenging (data not shown). At the same time, TF-iMG preserved multiple defining features of human microglia, including expression of canonical markers, ramified morphology, inflammatory responsiveness to IL1B and IL4, and phagocytic activity. These properties positioned TF-iMG as a useful compromise between biological relevance and experimental scalability, two features that are often difficult to achieve simultaneously in microglia-focused functional genomics.

### TF-iMG captured intrinsic and stimulus-induced microglial state dynamics

Our single-cell analyses further showed that TF-iMG had structured and biologically meaningful microglial states similar to previous findings [16, 43–47]. Even in the absence of stimulation, TF-iMG displayed homeostatic, activated, inflammatory, and proliferative states, with strong correspondence to human microglial states reported *in vivo*. This intrinsic heterogeneity was important because it suggested the model captured a degree of endogenous microglial plasticity instead of oversimplifying microglia into one uniform *in vitro* identity. Moreover, stimulation with IL1B and IL4 reproducibly generated separable pro-inflammatory and anti-inflammatory states, creating a tractable framework for studying microglial polarization and for anchoring genetic perturbation effects. The observation that these states were also associated with distinct Alzheimer’s disease risk genes further supported the disease relevance of the model.

### Arrayed Perturb-seq and multi-modal phenotyping enabled pathway validation and target identification

Another advancement was the establishment of an arrayed Perturb-seq workflow adapted to microglia target identification. In a proof-of-concept screen, perturbation of IL1B and IL4 pathway components shifted cells in the expected directions and attenuated stimulus-specific transcriptional responses. These findings validated the platform’s ability to recover known signaling relationships using single-cell readouts. Knockout of TGFBR2 demonstrated that we could detect broad microglial dysregulation across transcriptomic, secreted, morphological, and behavioral levels. Together, these experiments establish confidence that the platform can robustly connect gene perturbations to microglial state outcomes.

Importantly, the second study showed that the platform was not limited to validating expected pathway biology, but could also resolve more nuanced effects of disease-relevant regulators. Perturbations of IRF8, TREM2, ABI3, and HIF1A did not simply switch inflammatory states in a binary mode. Instead, they produced graded redistributions across microglial states, along with perturbation-specific downstream gene expression changes and orthogonal phenotypic consequences. In microglia, many therapeutically relevant genes may not act as binary gatekeepers, but rather tune the balance between homeostatic, inflammatory, protective, and disease-associated states. A multimodal single-cell approach was therefore valuable because it could capture these subtle but biologically meaningful shifts.

### Value of integrating transcriptomic, functional, and imaging readouts

In many screening settings, transcriptomic hits can be difficult to prioritize without functional corroboration [10, 31, 96, 97]. One strength of this study was the integration of multiple orthogonal readouts within the same screening framework. Transcriptomic profiling provided the highest-resolution view of microglial state identity and perturbation response, but cytokine measurements and live imaging added important functional context. In several cases, perturbation-induced gene expression changes were mirrored by altered secreted inflammatory signals or by measurable differences in morphology and movement. This convergence across modalities increased confidence in hit interpretation and would provide a richer basis for prioritizing targets for follow-up studies.

More broadly, the modular design of the platform makes it adaptable to diverse experimental questions. Additional stimuli, advanced cell culture conditions, and phenotypic outputs could be incorporated without fundamentally changing the workflow. This flexibility may be particularly valuable for modeling disease-relevant microenvironments and for comparing how candidate targets behave across multiple contexts.

## Limitations and future directions

This study has several limitations: TF-iMG is an *in vitro* model and therefore cannot fully reproduce the developmental, environmental, and regional complexity of human microglia *in vivo*. Some of the observed microglia states and distribution between states may be influenced by the TF-induced differentiation strategy or by culture conditions. In addition, the perturbation panels tested in this study were relatively focused and were primarily intended to establish and assess the platform performance with illustrative use cases. Broader application to a larger set of targets, alternative perturbation approaches such as CRISPRi or CRISPRa, and more complex disease-relevant stimuli and co-culture systems would be interesting next steps.

Further mechanistic work will also be needed to validate the downstream pathways suggested by the multimodal phenotypes reported here. For example, the trajectory defects observed following perturbations such as IRF8 and TREM2 suggest possible roles in chemotaxis or cytoskeletal regulation. These hypotheses would require dedicated follow-up studies with dedicated functional and cellular biological assays. Similarly, the disease associations inferred from DAM and IAM enrichment analyses provide a useful framework for transcriptional state interpretation but should be tested in additional and more comprehensive biological systems.

## Conclusions

Overall, this study established TF-iMG with arrayed multimodal Perturb-seq as a practical platform for microglia target identification. TF-iMG provided a rapid and CRISPR-compatible human microglia-like model, while the perturbation workflow enabled systematic mapping of how genetic perturbations reshaped microglial states across transcriptomic, cytokine, morphological, and behavioral dimensions. By capturing both expected pathway biology and more subtle disease-relevant perturbation effects, this platform offered a scalable approach for dissecting microglial state regulation and may support broader applications in neuroinflammation research and drug discovery.

## Author contributions

L.J., C.F. and J.H. conceived the research. L.J., M.L. and S.H. performed experimental work with guidance from J.H. and K.W.. L.J. performed the computational work with guidance from C.F., J.H. and K.W. J.L. supported the automation of time-lapse imaging quantification pipeline. Y.K. supported the next-generation sequencing. X.C., J.Y. and B.S. supported the TF-iMG flow cytometry experiments. All authors participated in interpretation of and in writing the manuscript.

## Acknowledgements

We thank Beata Henry, Seon Hye Cheon, Alan Ho, and Emily Deng for assistance with high-throughput imaging, tissue culture and flow cytometry. We also thank Daniel Ho, Michelle Duan and Thomas Perkins for guidance on transcriptomic data analysis. We acknowledge Gizem Rizki, Ajeet Singh, Neno Pokrajac, Ioana Soaita, Valerie Van Steenbergen, Joelle Dorskind, Rob Maher, Martin Henault, and Frederic Sigoillot for valuable discussions on CRISPR screening strategies. We are grateful to Robert Ihry, Stephan Reiling, and all members of Neuroscience Targets and Technologies and Data Science teams at Novartis Biomedical Research for insightful discussions and feedback on experimental design, data interpretation, and manuscript preparation.

## Methods

### Generation of TF-iMG hPSC line

We generated the TF-iMG hPSC line by introducing SPI1 and CEBPA transcription factors via piggybac (PB) system to a constitutively expressing Cas9 hPSC line previously established [32]. Cas9-expressing hPSC line were maintained in mTeSR1 (Stemcell Technologies, 85850) with 4ug/mL Blasticidin S HCl (Gibco A11139-03) in 6-well tissue culture plates (Falcon, 353046) coated with 1X hESC qualified Matrigel (Corning, 354277) diluted in DMEM/F12 (Gibco, 11320-033) prior to engineering. After reaching 85% confluency, hPSCs were switched to mTeSR1 containing 1uM Thiazovivin (Selleckchem, S1459) for two hours prior to being dissociated into single cells using pre-warmed Accutase (Gibco, A11105-1) for five minutes at 37°C. Cells were then pelleted at 300rcf for 3 minutes, resuspended in mTeSR1 with 1uM Thiazovivin and counted using the Beckman Coulter Vi-CellXR Cell Viability Analyzer. One million hPSCs were aliquoted into a 1.5mL Eppendorf tube and gently resuspended in transfection buffer containing 105uL Neon Resuspension Buffer R (Invitrogen, MPK10096R), 2ug of PB-SPI1-P2A-CEBPA plasmid, and 1ug of PB dual Helper plasmid. hPSCs were electroporated using the Neon Transfection System (Invitrogen, MPK5000) at 1050V, width 30, 2 pulses and immediately plated into two 10cm tissue culture dishes (Corning, 353003) containing mTeSR1 with 1uM Thiazovivin with approximately 500k cells per dish. The following day, a complete media change was performed with mTeSR1 containing 4ug/mL Blasticidin S HCl, 250ug/mL Geneticin (Gibco, 10131-035), and 1uM Thiazovivin. The second day following electroporation cells received another complete media change omitting Thiazovivin. All subsequent daily media changes were performed with mTeSR1 containing 4ug/mL Blasticidin S HCl and 250ug/mL Geneticin.

Following selection with Geneticin in 10cm dishes, clones were picked into 24 well plates (Corning, 3526) coated with 1X hESC qualified Matrigel and cultured in TeSR E8 (Stemcell Technologies, 05990) media containing 4ug/mL Blasticidin S HCl and 250ug/mL Geneticin. Colonies were expanded into 6-well plates using the same coating and media conditions and switched back to mTeSR1 containing 4ug/mL Blasticidin S HCl and 250ug/mL Geneticin following successful expansion. Cells were banked in CryoStor CS5 (Biolife Solutions, 205102) for storage. One TF-iMG clone was selected for all the subsequent experiments and studies.

GFP expressing TF-iMG line was established by transducing EF1a-GFP-puro lentivirus at the stem cell stage. Briefly, stem cells were dissociated to single cells with Accutase at 37C for 5 minutes. 180uL of lentiviruses were mixed with 400K cells in 820uL of mTeSR, and the mixture was plated in one well of a 6 well plated coated with Matrigel. The next day, change media containing 4ug/mL puromycin (Gibco, A1113803) and continue to maintain this culture as pool. GFP expressing TF-iMG cells were used in studies involved in time-lapse experiments.

### TF-iMG differentiation, cryopreservation, and general culture conditions

Prior to differentiation, TF-iMG hPSCs were maintained and expanded in mTeSR1 containing 4ug/mL Blasticidin S HCl and 250ug/mL Geneticin in 6-well tissue culture plates coated with 1X hESC qualified Matrigel. Upon optimal confluency (75%-85%) hPSCs were dissociated into a single cell suspension via pre-warmed Accutase for 5 minutes at 37C and pelleted at 300rcf for 3 minutes. The hPSC pellet was then resuspended in the maintenance medium with the addition of 1uM Thiazovivin and counted using the Beckman Coulter Vi-CellXR Cell Viability Analyzer. hPSCs were then seeded in the desired vessel coated with 1X hESC qualified Matrigel at a density of 250k cells per well for a 6-well plate, 4 million cells in a T150 (Corning, 430825), or 66 million cells in a 5-layer cell stack (Corning, 3313). The following day (Day 0) media was changed to “Microglia Complete” medium comprised of Neurobasal Media (Gibco, 21103-049), 0.5X B27 Supplement (Gibco, 12587), 1X N2 Supplement (Gibco, 17502-048), 1X Glutamax (Gibco, 35050-062), 1X MEM NEAA (Gibco, 11140-050), 1X Penicillin-Streptomycin (10,000U/mL; Gibco, 15140-163), 100ng/mL IL-34 (Peprotech, 200-34), 50ng/mL TGFb (Peprotech, 100-21), 10ng/mL NT3 (Peprotech, 450-03), 10ng/mL mCSF (Peprotech, 300-25), 10ng/mL GDNF (Peprotech, af-450-10), 10ng/mL BDNF (Peprotech, 450-02), and 2ug/mL Doxycycline (Sigma, D9891). TF-iMGs are mature and ready for experiments on Day 3.

For cryopreservation, three days after switching to “Microglia Complete” media (Day 3), TF-iMG were washed once with DPBS (Gibco, 14190-144) and dissociated by incubating in pre-warmed Accutase for 3 minutes at 37°C. Cells were pelleted at 300rcf for 5 minutes, resuspended in CryoStor CS10 (Biolife Solutions, 210102), and transferred to cryovials to bank. These cells are hereafter referred as d3 TF-iMG. For subsequent experiments, d3 TF-iMG were thawed and seeded on plates coated with 1X hESC qualified Matrigel diluted in DMEM/F12 and cultured in “Microglia Complete” media before use.

### TF-iMG Immunostaining and Imaging

d3 TF-iMG were thawed and seeded on a 96-well PhenoPlate (Revvity, 6055300) at a density of 30k cells/well in “Microglia Complete” media. Three days later, cells were fixed in 4% paraformaldehyde (PFA, Electron Microscopy Sciences, 15710) diluted in PBS (Gibco, 14040-133) for 15 minutes. PFA was removed and fixed cells were washed three times using PBS. Cells were blocked and permeabilized for 2 hours at room temperature in solution containing 10% Donkey Serum (Rockland, D120-00-0050) and 0.1% TritonX-100 (BioVision, 2124-50) in PBS. Primary antibodies were diluted in the blocking/permeabilization solution: Rabbit anti-human Iba1 (Fujifilm, 019-1974) was diluted 1:1000, mouse anti-human CD68 (BioRad, MCA5709) was diluted 1:500, and rabbit anti-human CX3CR1 (BioRad, AHP1589) was diluted 1:200. Cells were incubated with primary antibody overnight at 4C. The following day, primary antibody was washed out using PBS. Secondary antibodies Alexa Fluor Donkey anti-Rabbit 488 (Life Technologies A21206) and Alexa Fluor 594 Donkey anti-mouse (Life Technologies, A21203) were diluted 1:1000 in the blocking/permeabilization solution and DAPI (ThermoScientific, 62248) was diluted 1:250. Cells were incubated with secondary antibody and DAPI for 2 hours at room temperature. Secondary antibody was washed out with PBS. Cells were imaged on Opera Phenix Plus using a 40X objective.

### TF-iMG Phagocytosis Assay

d3 TF-iMG were thawed and seeded on a 96-well pre-coated PDL plate (Greiner, 655946) at a density of 30k cells/well in “Microglia Complete” media. Three days later, 50% of media was removed and replaced with “Microglia Complete” media containing 0.2uM Cell Tracker Green CMFDA Dye (Invitrogen, C7025), and 2ug/mL Hoechst 33342 (Promokine, PK-CA707-40046). Cells in the positive control condition had 20uM Cytochalasin D (Tocris, 1233) included the media. After incubation at 37°C for one hour cells were given a complete media change with “Microglia Complete” media containing pHrodo Red S. aureus Bioparticles conjugate (Invitrogen, A10010) and positive control cells retained 10uM Cytochalasin D in their media. Imaging commenced immediately using Opera Phenix with a 20X objective at one-hour intervals for a total of seventeen hours. Analysis of pHrodo intensity was conducted using Harmony 4.9 analysis software.

### TF-iMG Polarization and Cytokine Release Assay

For TF-iMG characterization, 30K/well of d3 TF-iMG were thawed and seeded on a 96-well pre-coated PDL plate (Greiner, 655946). Three days later, 50% of media was removed and replaced with “Microglia Complete” media containing 2X concentration of appropriate stimuli, resulting in a final concentration of 50ng/mL IL1b (R&D Systems, 201-LB/CF) or BSA control (Gibco, 15260037, 0.0001%) for 24 hours.

For IL1B/IL4/BSA-specific polarization experiments, 50K d3 TF-iMG were seeded on a 96-well pre-coated PDL plate (Greiner, 655946). Next day, IL1B (Peprotech, 200-01B-10UG, 50ng/ml), IL4 (Peprotech, 200-04-20UG, 50ng/ml) or BSA control (Gibco, 15260037, 0.0001%) were added to the media via half-media change. The supernatants were collected after 24 hours.

For scRNAseq profiling experiment stimulated with IL1B/IL4/ACM/BSA, 1 million of d3 TF-iMG were thawed and seeded with “Microglia Complete” media in a 6-well plate. On day 4, IL1B (Peprotech, 200-01B-10UG, 50ng/ml), IL4 (Peprotech, 200-04-20UG, 50ng/ml), ACM (ScienCell, 1811, 50%), or BSA control (Gibco, 15260037, 0.0001%) were added to the media via half-media change. Cells were harvest on day5 for scRNAseq described below.

For CRISPR editing experiments, 50K d3 TF-iMG were seeded on a 96-well pre-coated PDL plate (Greiner, 655946). Next day, half media was changed and lenti sgRNAs and Vpx-VLP particles were added. On day 6, we perform a full media change to remove lentivirus and Vpx-VLP. On day 8, IL1B/IL4/BSA polarization was practiced in the same way as the experiment above, and the supernatants were collected on day 9.

For these cytokine experiments, 80 uL was collected from each well of a 96-well plate and centrifuged at 800 rpm for 5 min. Approximately 60 ul of clarified supernatant was transferred and stored at −80°C until analysis. Secreted cytokines were quantified using the MSD V-PLEX Proinflammatory Panel 1 Human Kit (MSD, K15049D) and/or the R-PLEX Singleplex Human Osteopontin SPP1 Kit (MSD, F21YM-3), following the manufacturers’ standard protocols. Plates were read on a MESO QuickPlex SQ 600MM instrument. Three to four technical replicates in different wells were performed for each condition (Table S7).

### Ngn2 neuron differentiation and cryopreservation

We previously generated a constitutive Cas9-expressing line by targeting AAVS1 locus, followed by integration of PB-Ngn2 to create a doxycycline-inducible Ngn2 hPSC line [32, 98]. To differentiate these cells into Ngn2 neurons, hPSCs are seeded and expanded on appropriate cell culture surfaces coated with 1X hESC-qualified Matrigel at a density of 800,000 cells per 9.6 cmZ. Cells are suspended, seeded, and incubated in complete mTeSR1 medium supplemented with 1 µg/mL thiazovivin ROCK inhibitor

On day 0 differentiation, medium is fully replaced with fresh NGN2 medium without B-27^TM^ supplement (DMEM/F12, GlutaMAX medium, 1X MEM non-essential amino acids (NEAA), 1X Penicillin-Streptomycin (Pen/Strep), 1X N-2 supplement, 10ng/mL human NT-3 protein supplement, 10ng/mL human BDNF protein supplement and 2ug/mL doxycycline).

On day 2 differentiation, medium is fully replaced with fresh, complete NGN2-neuron medium (Sudhof media), including 1X B-27^TM^ supplement.

On day 3 differentiation, NGN2-neurons are dissociated from cell culture surface using Accutase for 5 to 10 minutes. Cells are centrifuged at 250g for 5 minutes and supernatant is removed. Pelleted cells are resuspended in CyroStor^®^ CS5 freeze medium (Stemcell Technologies, 07933) to appropriate cell density for long-term storage in liquid nitrogen. Temperature of resuspended cell tubes are gradually brought down using the VIA Freeze instrument to avoid crystallization of cell suspensions before being stored in liquid nitrogen.

### sgRNA plasmid design and lentivirus packaging

sgRNAs were designed using Broad Genetic Perturbation Platform CRISPick software (https://portals.broadinstitute.org/gppx/crispick/public). Human GRCh38 (NCBI), CRISPRko, and spyoCas9(NGG) options were chosen to pick sgRNAs. Top 3 ranked sgRNAs were cloned individually to lentigRNA plasmids containing either TagRFP or SNAP26f tag for transduction efficiency visualization. For any genetic KO experiments with lentigRNAs, the 3 individually packaged viruses per gene were mixed equally to maximize KO effect. All sgRNA sequences can be found in Table S6.

Lenti-X 293T cells (Takara, 632180) at passages 2–5 were recovered in penicillin/streptomycin-free DMEM (Gibco, 11995065) supplemented with 10% FBS and 1x NEAA (Gibco, 11140050) and maintained at 37°C. Cells were seeded onto collagen I-coated 6-well plates (Corning, 356400) at 0.75 million cells per well one day before transfection.

On the day of transfection, TransIT-293 reagent (Mirus, MIR 2700) and Opti-MEM serum-free medium (Invitrogen, 11058021) were warmed to room temperature and pre-mixed (16 ul TransIT-293 + 260 ul Opti-MEM per well). For each well, 2.78 ug sgRNA plasmid and 3 ug Cellecta Packaging Mix (Cellecta, CPC-K2A) were added to the transfection mixture. The master mix was incubated at room temperature for 15 min before being added to the cells. The following day, the medium was replaced with 1 mL fresh microglia Basal Medium (“Microglia Complete” media without all growth factors, DOX and Penicillin-Streptomycin). Viral supernatants were collected 72 h after transfection, passed through a 0.45 um cellulose acetate filter, and stored at −80°C. Vpx virus-like particles (VLPs) were packaged using Vpx containing plasmid and pMD2.G. Viral supernatant collection is the same as lenti sgRNA. Lentiviral titers were estimated based on the percentage of RFP-positive TF-iMG cells measured by flow cytometry 3 days after transduction.

### Vpx-assisted CRISPR editing in TF-iMG

Studies involving lentiviral sgRNA-mediated CRISPR editing were co-delivered with Vpx VLPs. Fresh or cryopreserved d3 TF-iMG cells were plated onto Matrigel-coated 6-well plates at 1.5 M cells per well or 96-well plates at 50 K cells per well and incubated overnight at 37°C. On day 4, a half-medium change was performed, followed by addition of lentivirus at MOI = 1 together with Vpx VLPs (1 uL per well for 96-well plates; 30 uL per well for 6-well plates). A full medium change was performed on day 6, followed by half-medium changes every 2 days thereafter.

### CRISPR editing efficiency assay (TF-iMG vs NGN2 Neuron)

To evaluate indel generation, TF-iMGs (DIV3) or NGN2 neurons (DIV3) were thawed from frozen stock. 1 million TF-iMGs or 500K NGN2 neurons per well were seeded onto Matrigel coated 24 well plates in respective culture media described above. On DIV4, 15ul Vpx and 15ul gRNA viral supernatant were added to the cells in a final culture volume of 250ul. On DIV6, 500ul media was added to the wells, and on DIV9 a full media change was performed. Thereafter, media was changed twice per week until the end of the experiment. On collection days (DIV10), media was removed, cells were washed once with DPBS, and genomic DNA was extracted using the DNeasy Blood and Tissue kit (Qiagen, 69504), in 50ul lysis buffer/well. Following extraction, genomic DNA was quantified on a nano-drop, and 50-100ng was used as template in a PCR reaction with primers designed to amplify the region targeted for CRISPR, and Herculase II fusion enzyme (Agilent, 600675), with the cycling conditions recommended by the manufacturer. PCR fragments were gel purified from a 0.8% agarose gel by spinning the gel slice through filter paper in a microfuge for 1 minute at 10,000 rpm. 50ng gel purified DNA was submitted for Sanger sequencing at Genewiz, with 5uM final concentration of sequencing primer/ reaction. Sequencing results were submitted to the TIDE NKI website (https://tide.nki.nl/) for indel analysis. All primer sequences can be found in Table S6.

### TF-iMG Time-lapse imaging

Thirty minutes before imaging, cells in 96-well plates were washed once with Neurobasal medium to remove dead cells and debris. Subsequently, 100 ul “Microglia Complete” media containing either 0.0001% BSA, 50 ng/mL IL1B, or 50 ng/mL IL4 was added to each well. Cells in the arrayed 96-well format were imaged using a CellVoyager CQ1 (Yokogawa) with a 10x objective in the GFP channel. Four fields per well were acquired every 10 min for 24 h. Four technical replicates in different wells were performed for each condition.

Single-cell morphology and trajectory features were quantified using TrackMate [99]. For cell detection and morphology measurements, the intensity threshold was set to 200. Cell tracking was performed using the Kalman tracker with the following parameters: initial search radius = 50, search radius = 50, and maximum frame gap = 5. All 19 default TrackMate measurements were collected. Only cells tracked for at least 40 frames were retained for downstream analysis (Table S8).

### Statistical tests

For single-measurement, two-group comparisons of cytokine release and imaging-based quantifications (for example, KO or treatment versus control), non-parametric Mann-Whitney U tests were used. For multi-group comparisons of imaging-based quantifications across multiple knockout conditions, one-way ANOVA followed by Dunn’s test was applied. GO Biological Process and KEGG pathway enrichment analyses were performed using ShinyGO 0.85 [100]. Only terms containing 40–500 genes were included, and redundant terms were removed (Table S3). For custom signature enrichment analyses, fold enrichment was assessed against a background gene list comprising genes expressed in at least 10 percent of cells across TF-iMG scRNA-seq datasets, and the statistical significance was calculated using Benjamini-Hochberg-adjusted p-values derived from Fisher’s exact test (Table S9).

### Flow cytometry

Microglia were detached in cold PBS, washed with FACS buffer (2% FBS in PBS), and adjusted 100-500 K cells per well. Cells were stained using the following antibodies: CD11b-PE/Cy7 (BioLegend, 301321) at 1:50, P2RY12-APC (BioLegend, 392119) at 1:50, and AlexaFluor-647 anti-CX3CR1 (BioLegend, 341607) at 1:200. Cells were incubated for 30 min at 4°C, washed twice with FACS buffer, and resuspended in 100 uL FACS buffer for flowcytometry analysis on Cytoflex (Beckman Coulter).

### Bulk RNA-seq library construction and data analysis

For bulk RNA-seq, 1 million day-5 TF-iMG cells cultured in a 6-well plate were detached and dissociated using Accutase (Stemcell Technologies, 07920) for 3 min at 37°C, followed by one wash with neurobasal medium (3 min, 300g). Total RNA was extracted using the RNeasy Plus Mini Kit (Qiagen, 74134). RNA-seq libraries were generated using the NEBNext Ultra II Directional RNA Library Prep Kit for Illumina (NEB, E7760S), and libraries were amplified for 9 PCR cycles. Following size selection using SPRIselect beads (Beckman, B23317) and library quantification with the NEBNext Library Quant Kit for Illumina (NEB, E7630L), final libraries at 150 pM were sequenced on a NovaSeq 6000 SP v1.5 100-cycle kit (Illumina, 20028401) using the following run configuration: R1 61, I1 8, I2 8, R2 61.

FASTQ files were aligned to GRCh38 and quantified using the exon quantification pipeline (EQP), an integrated alignment and counting workflow that outputs raw counts and TPM values [101] (Table S10). Spearman rank correlation analysis was then used to compare TF-iMG transcriptomes with published in vivo and in vitro microglia datasets [37].

### scRNA-seq library construction and data analysis

TF-iMG dissociation and washing for scRNA-seq were performed as described for bulk RNA-seq, except that cells were processed in Eppendorf LoBind 96-well Deepwell plates (Eppendorf, EP0030504305-20EA). Arrayed cell pellets were then fixed, combinatorially barcoded, and converted into indexed libraries using Evercode Cell Fixation v3 (Parse Biosciences, ECFC3300) and Evercode Mini WT v3 kits (Parse Biosciences, ECWT3100 and ECWT3300) according to the manufacturers’ standard protocols. cDNA was amplified for 9 cycles and final libraries were amplified for 8 cycles. After size selection using SPRIselect beads (Beckman, B23317) and library quantification using the NEBNext Library Quant Kit for Illumina (NEB, E7630L), final libraries at 150 pM were sequenced using NovaSeq X Series 10B 100-cycle kits (Illumina, 20085596) with the following run configuration: R1 64, I1 8, I2 8, R2 58.

Sub-library FASTQ files were demultiplexed by sample, aligned to GRCh38, converted to count matrices, and integrated across sub-libraries and samples using the split-pipe v1.4 pipeline[49]. For lentiviral marker detection, a custom GRCh38_LV2 reference was generated by appending the co-transcribed RFP and PuroR sequences from the sgRNA delivery lentiviral construct to the gene annotation file as a custom gene “LentiMarker” (Table S11).

The integrated raw count matrix was analyzed in Seurat v5 using R 4.6.0 [102]. Cells were filtered using the criteria: nFeature_RNA < 15000, nCount_RNA < 100000, and percent.mt < 15. Data were log-normalized and scaled, and the top 2,000 variable features were used for principal component analysis and clustering at resolution = 0.2. UMAP was used for dimensionality reduction and visualization. For pseudotime trajectory analysis, the Seurat object was converted to a CellDataSet (cds) object compatible with Monocle 3, and single-cell trajectories were inferred via the core function “cell_graph [50, 51]. Pseudotime value for each cell was assigned by assigning the “homeostatic” node as the start state via function “order_cell”, and the complete pseudotime trajectory was embedded in the original UMAP via function “plot_cells” [50, 51]. For differentially expressed genes (Table S2 and S4), cluster-specific or KO-condition-exclusive marker genes were identified using the FindAllMarkers function, with significance determined using the default Wilcoxon rank-sum test. Differentially expressed genes between KO and control conditions were identified using the FindMarkers function with the MAST framework specified by test.use = “MAST” [103]. The default log2 fold-change cutoff was set to 0.5 except for DEG or marker inputs for pathway term enrichment which was set to 0.25 against the background or control conditions.

## Key Resources Table

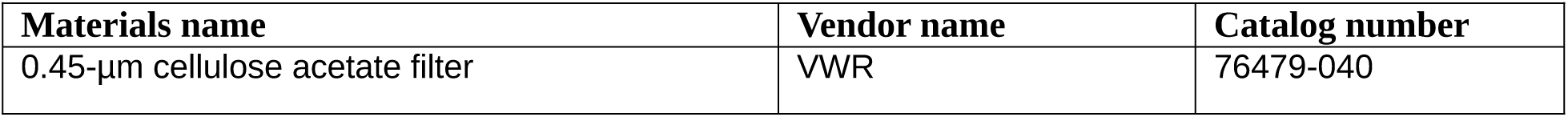
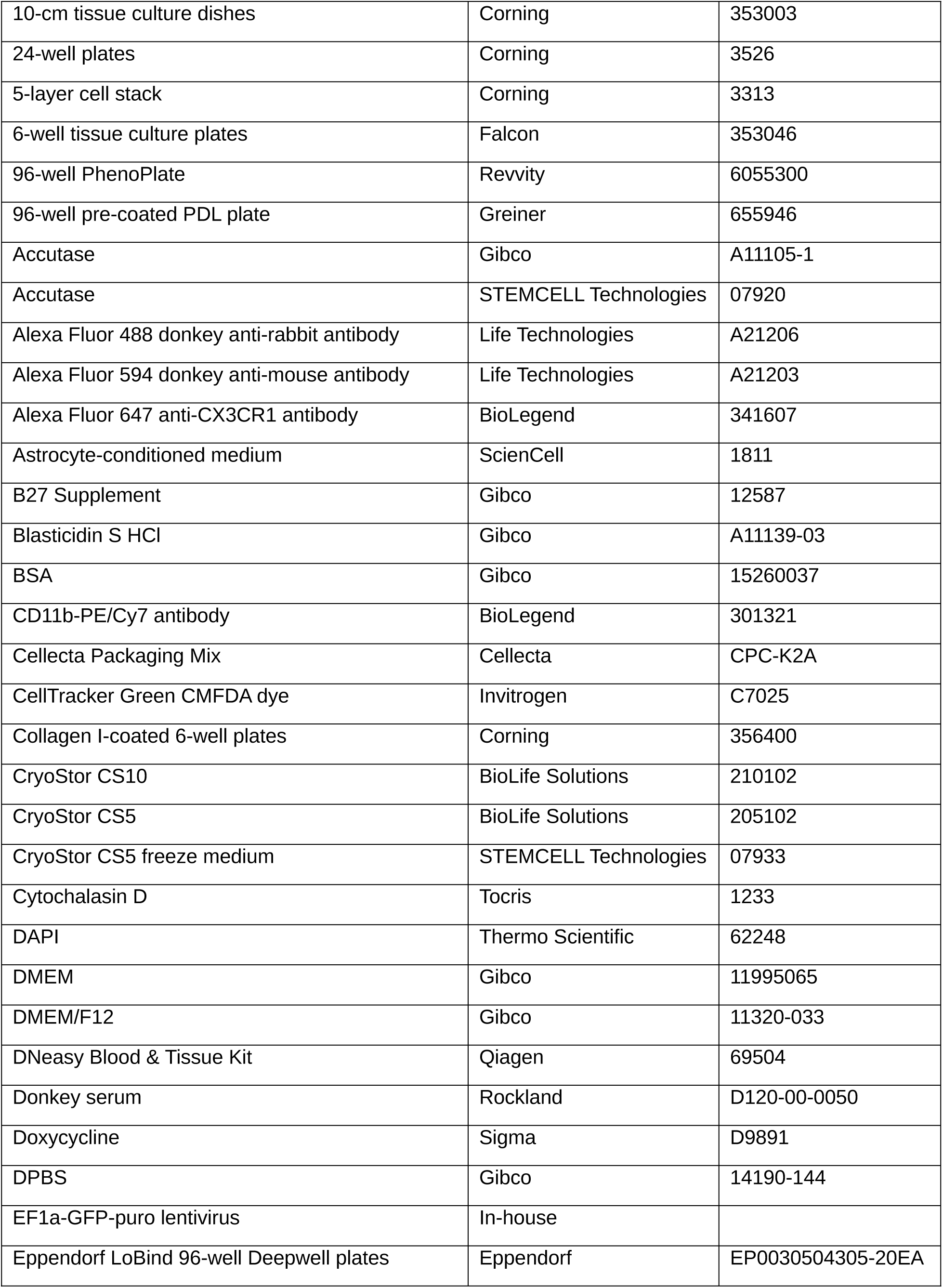
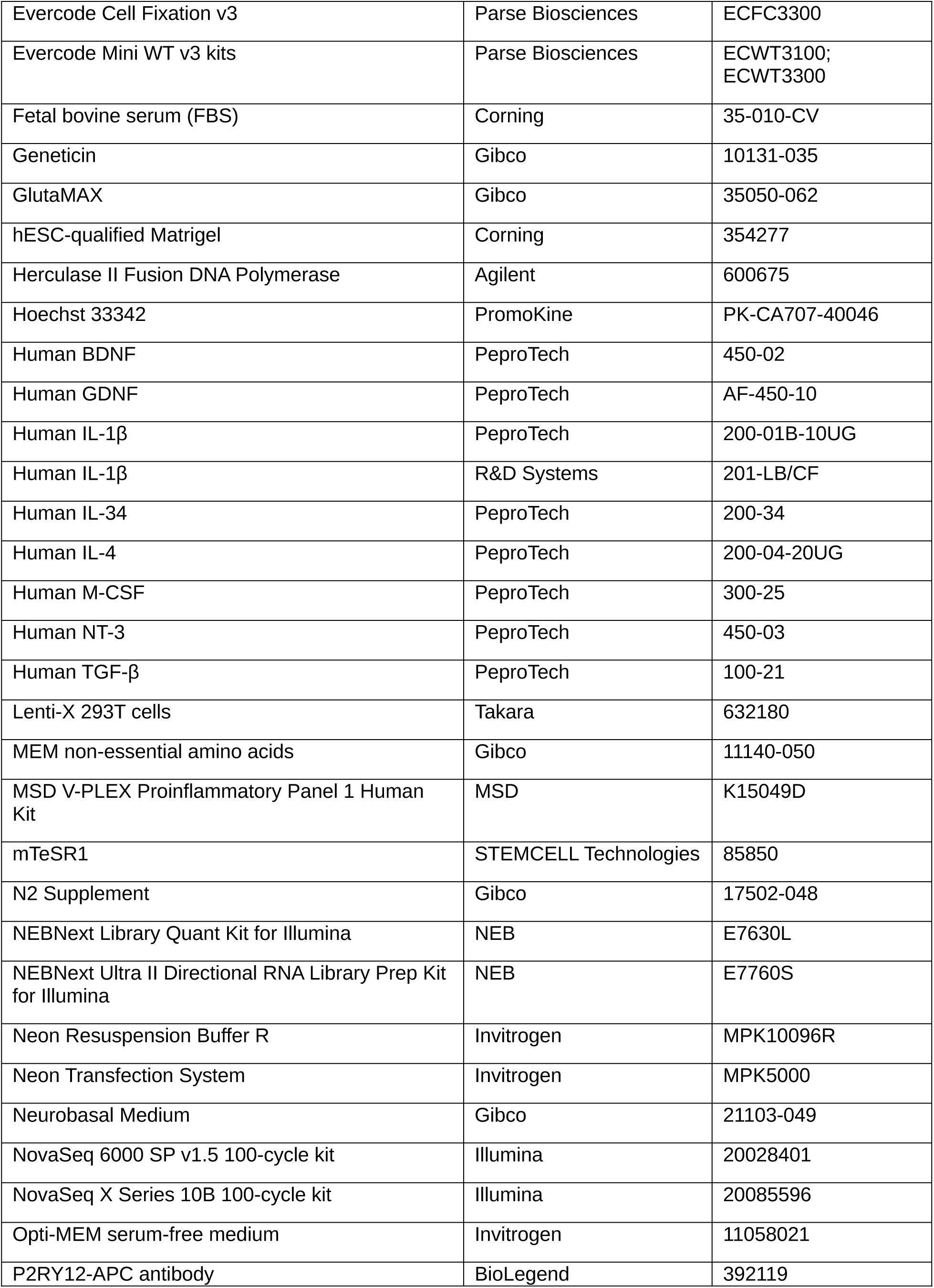
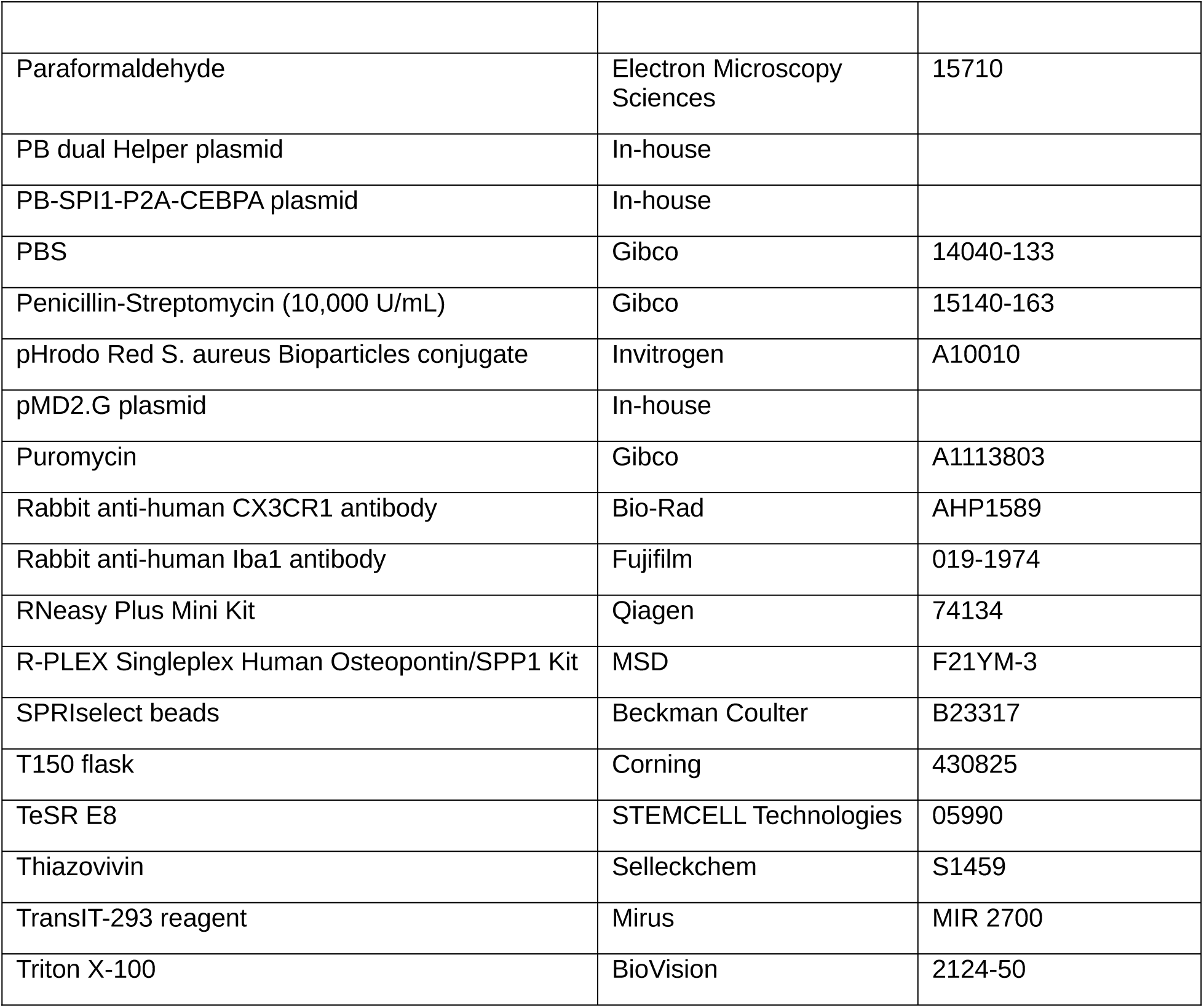

## Supplementary Tables

**Table S1:** Cell numbers by state and condition (treatment or treatment * perturbation) across all TF-iMG scRNA-seq datasets

**Table S2:** Cluster marker genes for all TF-iMG scRNA-seq datasets

**Table S3:** Full list of KEGG or GO Biological Process enrichments

**Table S4:** Condition-associated (KO vs Ctrl, or KO-specific) DEGs across all TF-iMG scRNA-seq datasets

**Table S5:** DAM and IAM signature gene sets

**Table S6:** sgRNA and primer sequences used for CRISPR knockout and editing efficiency assay in TF-iMG

**Table S7:** Source data for TF-iMG cytokine release measurements

**Table S8:** Source data for TF-iMG morphology and trajectory quantitative measurements from time-lapse imaging

**Table S9:** Background gene list comprising genes expressed in at least 10 percent of cells across TF-iMG scRNA-seq datasets

**Table S10:** Bulk RNA-seq TPM values for TF-iMG and other previously published microglia models

**Table S11**: Sequence of custom gene “LentiMarker”

## Data Availability

Raw sequencing data and processed gene-expression data that support the findings of this study have been deposited in NCBI Gene Expression Omnibus (GEO) under SuperSeries accession GSE354700, comprising bulk RNA-seq (GSE354659) and single-cell RNA-seq (GSE354698). Raw imaging data and scRNA-seq Seurat R objects are available at Zenodo (https://zenodo.org/records/20615641). All other source data are provided in supplementary tables with this paper.

