## Supplementary Figures S1-S4 and legends for "A multimodal arrayed Perturb-seq platform for dissecting microglial state dynamics and target identification"

Fig.S1

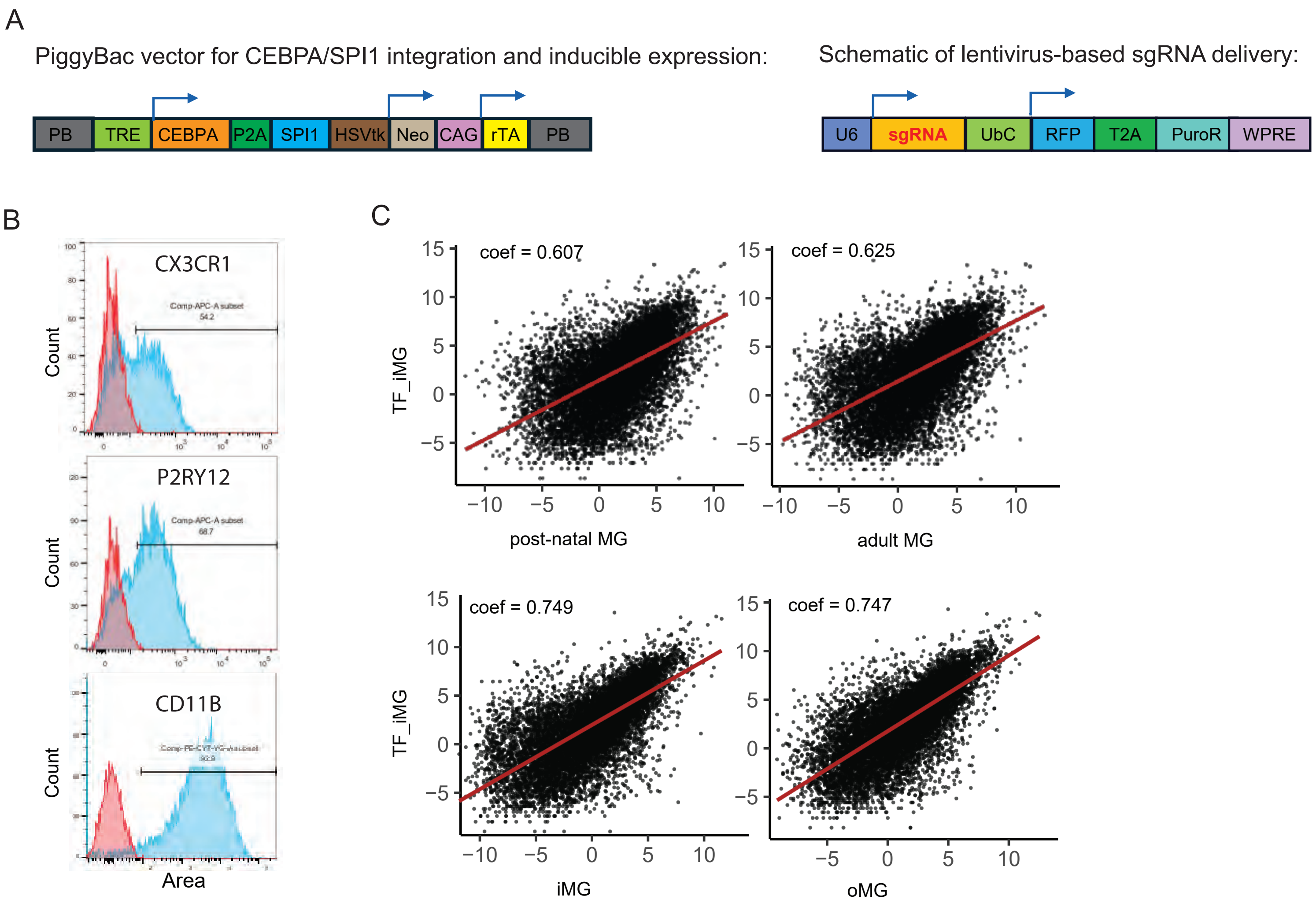

A
PiggyBac vector for CEBPA/SPI1 integration and inducible expression:
PB
TRE
CEBPA
P2A
SPI1
HSVtk
Neo
CAG
rTA
PB
Schematic of lentivirus-based sgRNA delivery:
U6
sgRNA
UbC
RFP
T2A
PuroR
WPRE
B
CX3CR1
P2RY12
CD11B
Count
Area
C
coef = 0.607
coef = 0.625
coef = 0.749
coef = 0.747
TF_iMG
post-natal MG
adult MG
iMG
oMG

#### Fig. S1

Fig. S1A. Schematic of vectors. Left, PiggyBac vector for CEBPA/SPI1 integration and inducible expression; Right, lentivirus-based sgRNA delivery construct.

**Fig. S1B.** Flow cytometry histograms showing expression of microglial markers (CD11B, CX3CR1, and P2RY12) before and after doxycycline induction.

**Fig. S1C.** Scatter plots showing significant positive Spearman correlations between TF-iMG transcriptomes and published datasets from human postnatal microglia, adult microglia, hPSC-derived microglia-like cells (iMG), and their integrations into cerebral organoids (oMG).

Fig.S2

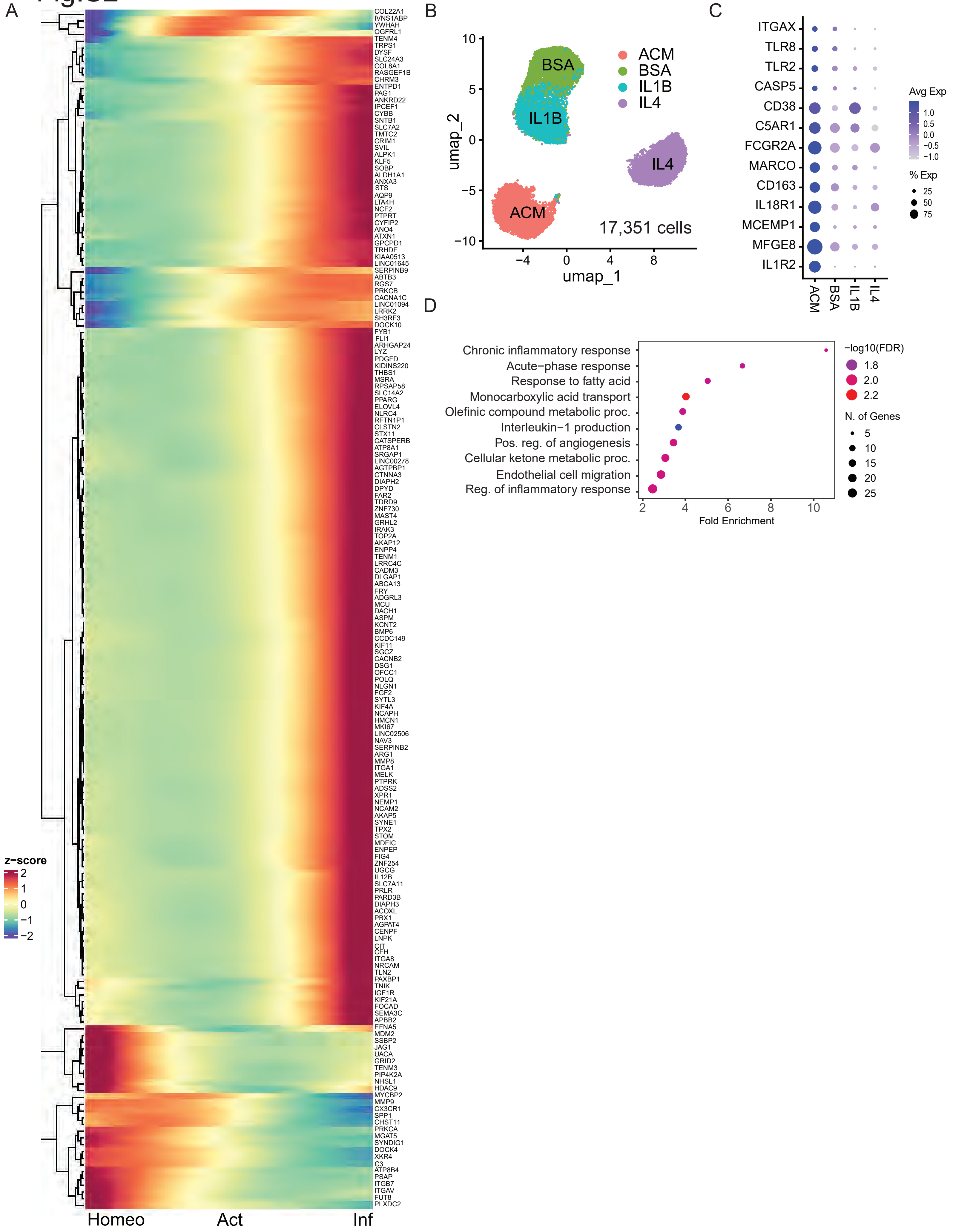
A
COL22A1
IVNS1ABP
YWHAH
OGFRL1
TENM4
TRPS1
DYSF
SLC24A3
COL8A1
RASGEF1B
CHRM3
ENTPD1
PAG1
ANKRD22
IPCEF1
CYBB
SNTB1
SLC7A2
TMTC2
CRIM1
SVIL
ALPK1
KLF5
SOBP
ALDH1A1
ANXA3
STS
AQP9
LTA4H
NCF2
PTPRT
CYFIP2
ANO4
ATXN1
GPCPD1
TRHDE
KIAA0513
LINC01645
SERPINB9
ABTB3
RGS7
PRKCB
CACNA1C
LINC01094
LRRK2
SH3RF3
DOCK10
FYB1
FLI1
ARHGAP24
LYZ
PDGFD
KIDINS220
THBS1
MSRA
RPSAP58
SLC14A2
PPARG
ELOVL4
NLRC4
RFTN1P1
CLSTN2
STX11
CATSPERB
ATP8A1
SRGAP1
LINC00278
AGTPBP1
CTNNA3
DIAPH2
DPYD
FAR2
TDRD9
ZNF730
MAST4
GRHL2
IRAK3
TOP2A
AKAP12
ENPP4
TENM1
LRRC4C
CADM3
DLGAP1
ABCA13
FRY
ADGRL3
MCU
DACH1
ASPM
KCNT2
BMP6
CCDC149
KIF11
SGCZ
CACNB2
DSG1
OFCC1
POLQ
NLGN1
FGF2
SYTL3
KIF4A
NCAPH
HMCN1
MKI67
LINC02506
NAV3
SERPINB2
ARG1
MMP8
ITGA1
MELK
PTPRK
ADSS2
XPR1
NEMP1
NCAM2
AKAP5
SYNE1
TPX2
STOM
MDFIC
ENPEP
FIG4
ZNF254
UGCG
IL12B
SLC7A11
PRLR
PARD3B
DIAPH3
ACOXL
PBX1
AGPAT4
CENPF
LNPK
CIT
CFH
ITGA8
NRCAM
TLN2
PAXBP1
TNIK
IGF1R
KIF21A
FOCAD
SEMA3C
APBB2
EFNA5
MDM2
SSBP2
JAG1
UACA
GRID2
TENM3
PIP4K2A
NHSL1
HDAC9
MYCBP2
MMP9
CX3CR1
SPP1
CHST11
PRKCA
MGAT5
SYNDIG1
DOCK4
XKR4
C3
ATP8B4
PSAP
ITGB7
ITGAV
FUT8
PLXDC2
z−score
2
1
0
−1
−2
Homeo
Act
Inf
B
10
5
0
−5
−10
umap_2
−4
0
4
8
umap_1
BSA
IL1B
IL4
ACM
ACM
BSA
IL1B
IL4
17,351 cells
C
ITGAX
TLR8
TLR2
CASP5
CD38
C5AR1
FCGR2A
MARCO
CD163
IL18R1
MCEMP1
MFGE8
IL1R2
ACM
BSA
IL1B
IL4
Avg Exp
1.0
0.5
0.0
−0.5
−1.0
% Exp
25
50
75
D
Chronic inflammatory response
Acute−phase response
Response to fatty acid
Monocarboxylic acid transport
Olefinic compound metabolic proc.
Interleukin−1 production
Pos. reg. of angiogenesis
Cellular ketone metabolic proc.
Endothelial cell migration
Reg. of inflammatory response
2
4
6
8
10
Fold Enrichment
−log10(FDR)
1.8
2.0
2.2
N. of Genes
5
10
15
20
25

#### Fig. S2

**Fig. S2A.** Heatmap of the top 200 genes showing dynamic expression along the homeostatic-to-inflammatory pseudotime trajectory in TF-iMG (z-scores).

**Fig. S2B.** UMAP of day-5 TF-iMG profiled 24 h after treatment with BSA, IL1B, IL4, or ACM.

**Fig. S2C.** Dot plot of top marker genes selectively enriched in TF-iMG after ACM treatment.

**Fig. S2D.** Dot plot of the top 10 GO Biological Process terms enriched among genes upregulated after ACM treatment in TF-iMG.

Fig.S3

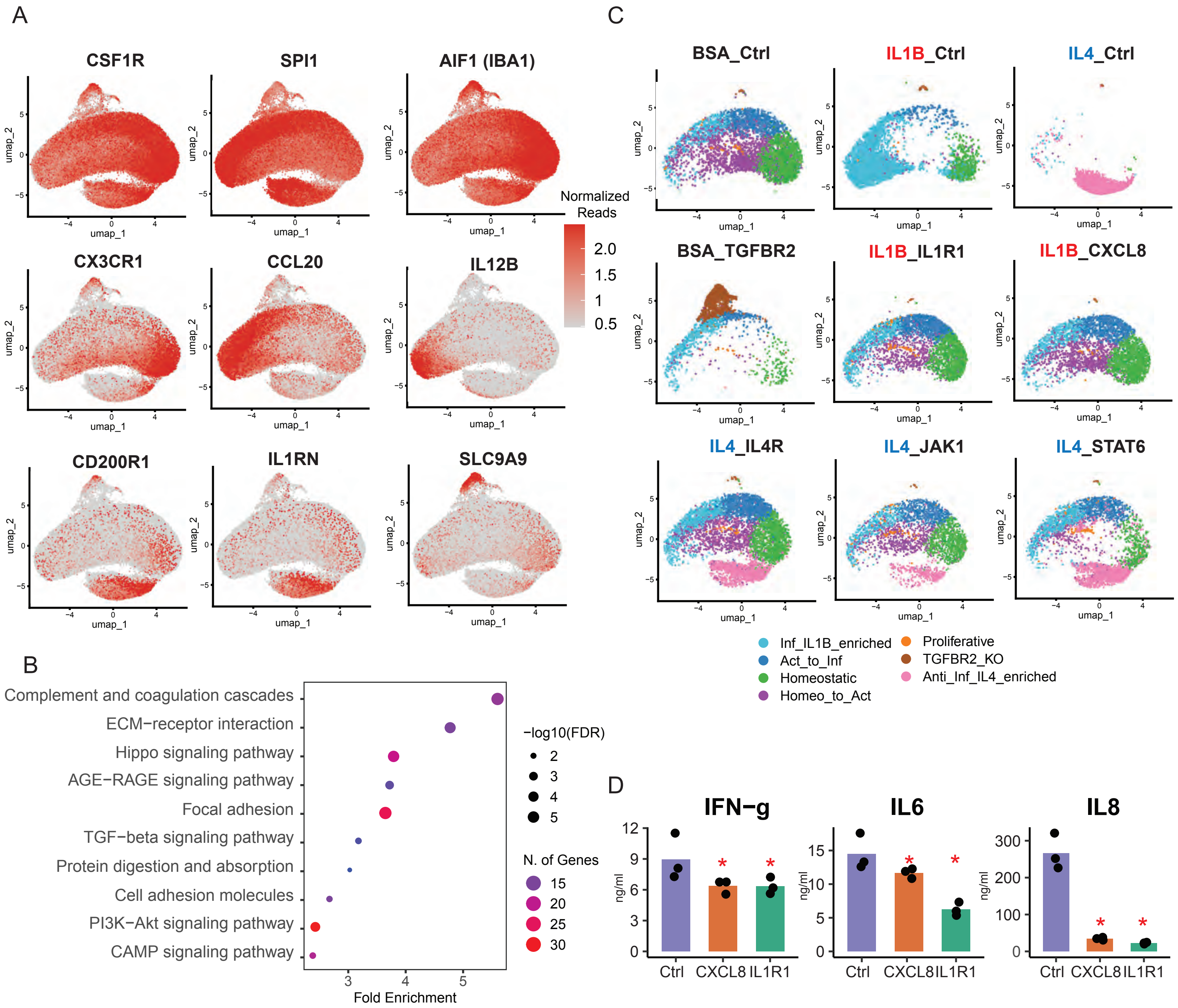
A
CSF1R
SPI1
AIF1 (IBA1)
CX3CR1
CCL20
IL12B
CD200R1
IL1RN
SLC9A9
Normalized Reads
2.0
1.5
1
0.5
umap_1
umap_2
B
Complement and coagulation cascades
ECM−receptor interaction
Hippo signaling pathway
AGE−RAGE signaling pathway
Focal adhesion
TGF−beta signaling pathway
Protein digestion and absorption
Cell adhesion molecules
PI3K−Akt signaling pathway
CAMP signaling pathway
Fold Enrichment
−log10(FDR)
N. of Genes
C
BSA_Ctrl
IL1B_Ctrl
IL4_Ctrl
BSA_TGFBR2
IL1B_IL1R1
IL1B_CXCL8
IL4_IL4R
IL4_JAK1
IL4_STAT6
Inf_IL1B_enriched
Act_to_Inf
Homeostatic
Homeo_to_Act
Proliferative
TGFBR2_KO
Anti_Inf_IL4_enriched
D
IFN−g
IL6
IL8
ng/ml
Ctrl
CXCL8
IL1R1

**Fig. S3**

**Fig. S3A.** Feature plots showing expression of pan-microglia and state-associated marker genes in the TF-iMG IL1B and IL4 pathway Perturb-seq dataset. Pan-microglia markers: CSF1R, SPI1, AIF1; homeostatic: CX3CR1; pro-inflammatory: CCL20, IL12B; anti-inflammatory: CD200R1, IL1RN; TGFBR2 knockout-associated: SLC9A9.

**Fig. S3B.** Dot plot of the top 10 KEGG pathways enriched among genes upregulated in the TGFBR2 knockout cluster.

**Fig. S3C.** UMAP in Fig.3B split by selected “treatment_perturbation” conditions.

**Fig. S3D.** Bar plots showing significantly reduced IFN-gamma (left), IL6 (middle), and IL8 (right) release following IL1B pathway perturbation (CXCL8 or IL1R1 knockout) compared with non-targeting control knockout under IL1B polarization. N = 3, $*p < 0.1$, Mann-Whitney U test.

### Fig.S4

A

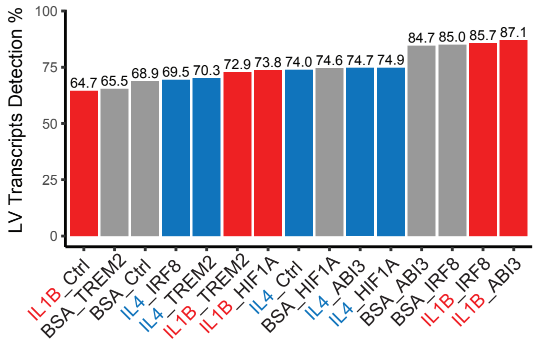


B

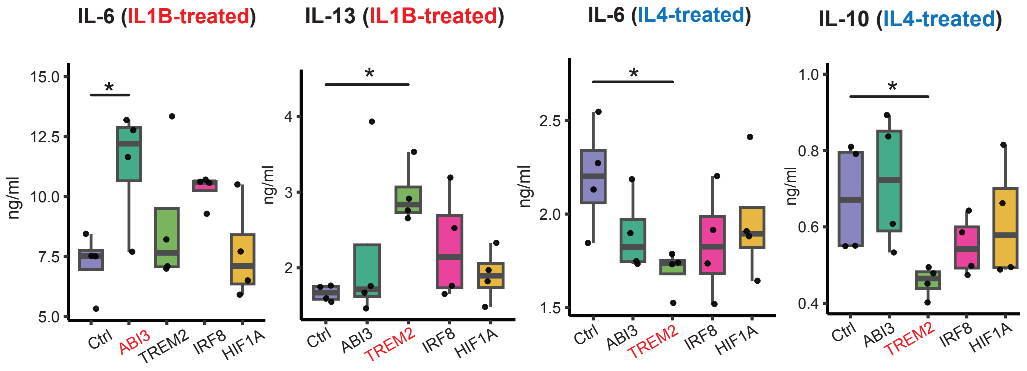


C

| Cytokines \ KOs | | ABI3 | TREM2 | IRF8 | HIF1A |
|---|---|---|---|---|---|
| IL1B-treated | IL-6 | UP | | DOWN | |
| | IL-8 | | | UP | DOWN |
| | IL-12 | | UP | UP | UP |
| | IL-13 | | UP | | |
| IL4 -treated | IL-6 | | DOWN | | |
| | IL-10 | | DOWN | DOWN | |

D

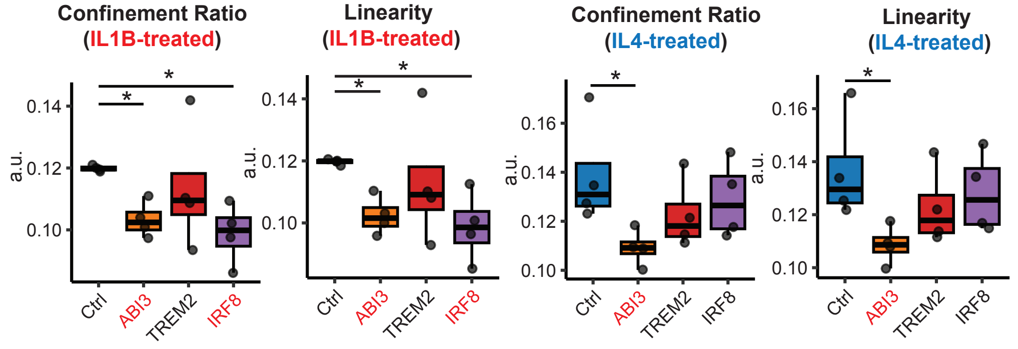


E

| Phenotypes \ KOs | | ABI3 | TREM2 | IRF8 |
|---|---|---|---|---|
| IL1B-treated | Linearity | DOWN | | DOWN |
| | Confinement ratio | DOWN | | DOWN |
| | Displacement | | | DOWN |
| IL4-treated | Linearity | DOWN | | |
| | Confinement ratio | DOWN | | |
| | Perimeter | | DOWN | |
| | Area | | DOWN | |
| | Shape Index | | DOWN | |

#### Fig. S4

**Fig. S4A.** Bar plots showing the percentage of cells with detectable lentiviral marker transcripts across the 15 conditions (red: IL1B-treated, blue: IL4-treated, gray: BSA-treated).

**Fig. S4B.** Extended box plots of significantly altered cytokine release under IL1B or IL4 polarization corresponding to Fig. 4H.

**Fig. S4C.** Summary of all significant cytokine changes shown in Fig. 4H and Fig. S4B.

**Fig. S4D.** Extended box plots of significantly altered cell morphology and trajectory parameters under IL1B or IL4 polarization corresponding to Fig. 4I.

**Fig. S4E.** Summary of all significant cell trajectory and morphology changes shown in Fig. 4I and Fig. S4D.
